# 2-Hydroxyglutarate Redirects Fatty Acid Partitioning to Mitigate Lipotoxic Stress and Preserve Metabolic Fuel

**DOI:** 10.64898/2026.08.12.744465

**Authors:** Niv Vigder, Amit Chandra, Nishith Shrimali, Sergey Tumanov, Vlad Elgart, Huamei He, Ryan Mulhern, Ram P. Chakrabarty, Navdeep S. Chandel, Stuart Cordwell, Steven P. Gygi, João A. Paulo, Joseph Loscalzo

## Abstract

The role of 2-hydroxyglutarate in lipid metabolism is currently unknown. Here we show that 2HG redistributes the partitioning of fatty acids into triglyceride storage and away from membrane phospholipid synthesis, mitochondrial oxidation, and lipotoxic intermediates. In primary human cardiac and vascular cells, both enantiomers, D2HG and L2HG, expanded triglyceride stores and lipid droplets while selectively depleting phosphatidylethanolamine, with L2HG acting more potently than D2HG despite lower intracellular accumulation. Mechanistically, L2HG increases DGAT-dependent triglyceride synthesis, slows triglyceride turnover, and constrains the ethanolamine branch of the Kennedy pathway. This response limits fatty acid oxidation, long-chain acylcarnitine accumulation, and lipid peroxidation independently of pseudohypoxic transcription or canonical lipid storage regulators, while also remodeling the phosphoproteome and redox proteome. L2HG accumulation induces hypertriglyceridemia in mice, redistributes the acyl chain composition of cardiac triglycerides, and limits ischemia-induced acylcarnitine accumulation in the heart, mirroring a positive association between circulating 2HG and triglycerides in humans. Thus, 2HG expands metabolic flexibility from whether fatty acids are used as fuel to how that fuel is allocated among storage, membrane synthesis, and oxidation.

## Introduction

Cells must continuously buffer the free fatty acids they take up and synthesize. In excess, free fatty acids and their metabolic derivatives, including acylcarnitines, diacylglycerol, ceramide, and oxidizable polyunsaturated phospholipids, become lipotoxic, damaging the endoplasmic reticulum and mitochondria and promoting membrane lipid peroxidation^1,2^. A principal defense against this stress is esterification of free fatty acids into inert neutral triglycerides (triacylglycerols) within lipid droplets, which sequester reactive lipid species away from membranes and oxidative machinery^3,4^. Accordingly, triglyceride storage protects mitochondria from fatty acid overload and shields polyunsaturated acyl chains from peroxidation^5,6^. The rate and extent to which fatty acids are partitioned into storage, rather than membrane synthesis or oxidation, therefore, represent a core axis of metabolic flexibility and help determine cellular vulnerability to lipotoxic injury^7^.

How this fatty acyl partitioning is controlled remains incompletely understood. Lipid droplet biogenesis is shaped by nutrient supply, transcriptional regulators such as sterol regulatory element-binding protein(s) (SREBP(s)), and lipid-droplet protein machinery^8^. However, whether a diffusible intracellular metabolite can itself set fatty acid fate, biasing acyl flux toward storage and away from membrane synthesis and oxidation, has not been established. Such a signal would provide a direct means to couple cellular redox, metabolic, and energetic state to lipid handling.

This mode of lipidomic control sits outside the canonical view of 2-hydroxyglutarate (2HG), a chiral metabolite best known in its *R*-form as an oncometabolite and epigenetic regulator and in its *S*-form as a regulator of central carbon metabolism in hypoxia or pseudo-hypoxic states^9,10^. The D-enantiomer (D2HG), produced by the neomorphic activity of mutant isocitrate dehydrogenase, accumulates to millimolar concentrations in glioma and leukemia cells where it competitively inhibits α-ketoglutarate-dependent dioxygenases, including TET DNA hydroxylases and Jumonji-domain histone demethylases, to remodel chromatin and block differentiation^11–15^. The L-enantiomer (L2HG) arises (patho)physiologically rather than by mutation, generated through promiscuous lactate- and malate-dehydrogenase activity under hypoxia^16,17^, acidosis^18^, and a reduced mitochondrial redox state^19^. Although long regarded as a low-abundance by- product of this enzymatic promiscuity, L2HG is increasingly recognized as a regulated physiological signaling metabolite with defined targets and measurable biological effects^19^. Like D2HG, L2HG can influence chromatin and signaling, reprogramming gene expression in renal cancer and lymphocytes and modulating HIF activity^19–21^, with its abundance constrained in part by the dedicated catabolic enzyme L2HGDH^22^. Across these contexts, both enantiomers have been viewed largely through the lens of α-ketoglutarate antagonism and epigenome regulation. More recently, mass spectrometry-based approaches have begun to identify covalent^23^ and non-covalent^24^ 2HG-binding proteins beyond canonical α-ketoglutarate-dependent dioxygenases, suggesting a broader target landscape.

Here, we show that 2HG redirects fatty acid fate. We found that both enantiomers, with L2HG acting more potently, shift fatty acids into stored triglycerides and away from membrane phospholipid synthesis and mitochondrial oxidation. This effect occurs through coordinated acceleration of triglyceride synthesis and slowing of triglyceride turnover, expanding triglyceride stores and lipid droplets. In parallel, 2HG selectively depletes phosphatidylethanolamine while sparing phosphatidylcholine, lowers mitochondrially lipotoxic long-chain acylcarnitines associated with incomplete β-oxidation, and decreases lipid peroxidation. This lipotype required the committed DGAT step of triglyceride synthesis, but not the pseudohypoxic transcriptional program or canonical lipogenic regulators induced by L2HG. Mice unable to degrade L2HG because of a genetic loss of functional L2HGDH develop hypertriglyceridemia, mirroring the positive association between circulating 2HG and triglycerides in human cohorts, and accumulate fewer long-chain acylcarnitines during myocardial ischemia. These findings establish 2HG as a regulatory metabolite of fatty acid fate, extending the reach of this central-carbon metabolite from epigenetic regulation to the flux-level partitioning of a macronutrient. More broadly, these findings expand the notion of metabolic flexibility beyond fuel choice, defining the allocation of a single fuel among several fates as a regulated metabolic decision.

## Results

### 2HG enantiomers differentially expand triglyceride stores in primary human cells

To test whether 2HG enantiomers differentially alter the lipotype of primary human cells, we treated primary human cells for 24 h with cell-permeant L2HG or D2HG octyl esters (OL2HG and OD2HG). Intracellular total O2HG accumulated similarly with both esters and was undetectable in vehicle-treated cells (Extended Data Fig. 1a), whereas free total 2HG increased ∼10-fold with OL2HG and ∼60-fold with OD2HG (Extended Data Fig. 1b). Chiral LC–MS confirmed enantiomeric fidelity: OL2HG-treated cells accumulated only L2HG (∼7 nmol mg^-1^ protein), whereas OD2HG-treated cells accumulated only D2HG (∼150 nmol mg^-1^ protein) (Extended Data Fig. 1c–e).

We next used non-targeted LC–MS/MS lipidomics to define the lipid response to 2HG. TGs were the dominant lipid class increased by 2HG. In cardiomyocytes, OL2HG increased 39 of 41 detected TG species by 2.9- to 9.0-fold (Fig. 1a), whereas OD2HG shifted the same species in the same direction more modestly (2.3- to 3.9-fold; Fig. 1b and Supplementary Table 1). Both enantiomers increased total cellular TG relative to DMSO (*P* < 0.0001 for each pairwise comparison), but OL2HG did so significantly more strongly (Fig. 1c), despite accumulating to ∼20-fold lower levels than OD2HG (Extended Data Fig. 1c–e). Thus, both enantiomers converge on TG accumulation, with greater apparent potency of OL2HG.

**Fig. 1:**
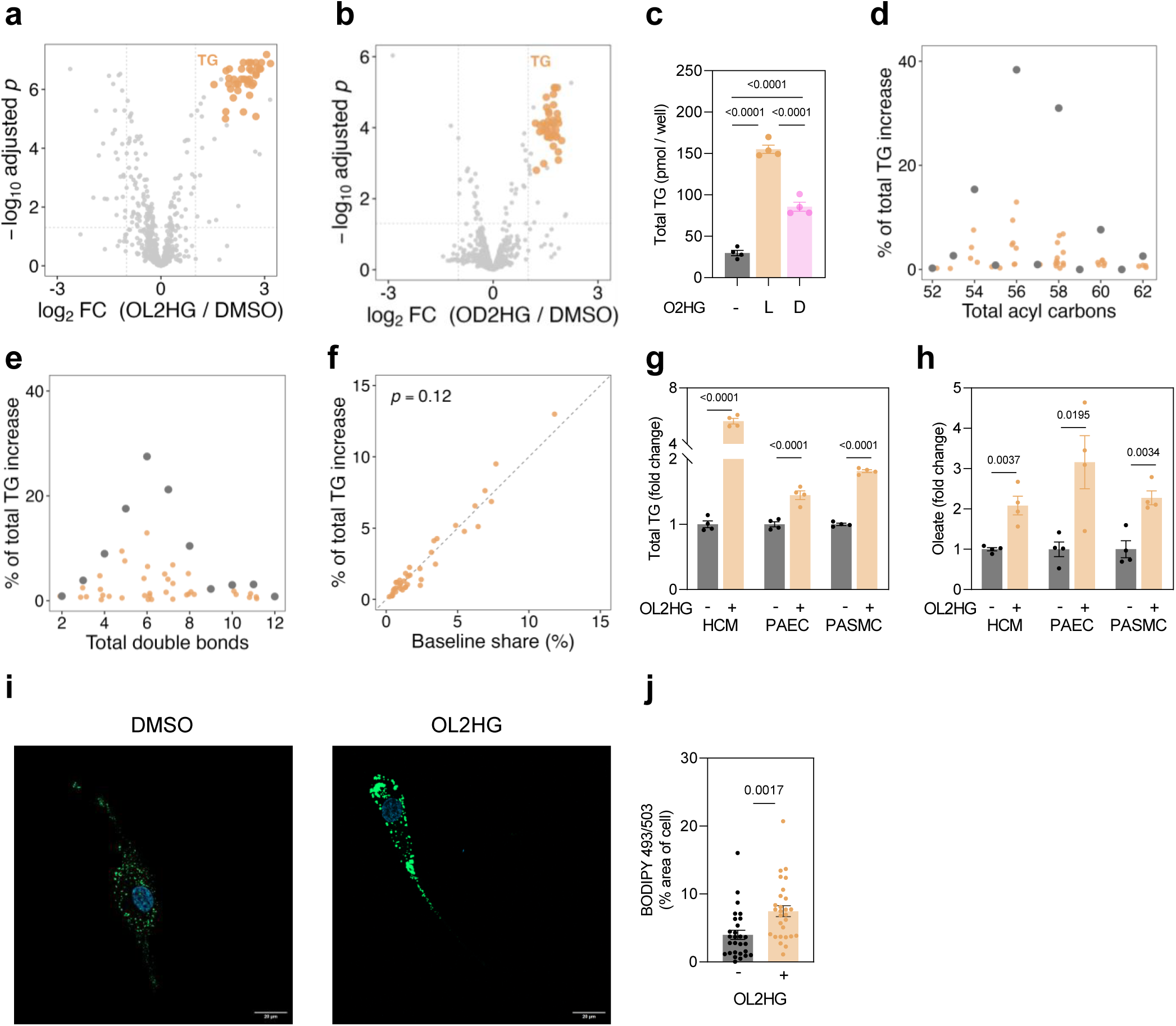
2HG enantiomers differentially expand triglyceride stores and lipid droplets in primary human cells. **a,b,** Lipidomic volcano plots from HCMs treated with DMSO, OL2HG (500 µM) (**a**), or OD2HG (500 µM) (**b**) for 24 h. TG species are orange; other lipids are grey. Significance: |FC| ≥ 2 and BH-adjusted *P* < 0.05; *n* = 4 per group. **c,** Total TG abundance in HCMs treated with OL2HG (500 µM) or OD2HG (500 µM) for 24 h; *n* = 4. **d,e,** Contribution of OL2HG-induced TG species to the total TG increase, grouped by acyl-carbon number (**d**) or double-bond number (**e**); orange, individual TG species; grey, summed bin. **f,** Contribution of each OL2HG-responsive TG species to newly accumulated TG plotted against its baseline pool share. Dashed line, y = x; slope ≈ 1.06, *P* = 0.12 by two-sided t-test of the fitted regression slope against 1, indicating proportional expansion. **g,h,** Total TG (**g**) and free oleate (**h**) in HCMs, PAECs and PASMCs treated with OL2HG (500 µM) for 24 h; *n* = 4 per group. **i,j,** Representative 63× confocal images (**i**) and quantification (**j**) of BODIPY 493/503-stained lipid droplets in oleate (50 µM)-loaded HCMs treated with DMSO or OL2HG (500 µM) for 24 h. Nuclei are stained with Hoechst. Scale bar, 20 µm; *n* = 28 cells from 4 independent samples per group. Unless otherwise indicated, data are mean ± s.e.m.. For two-group comparisons, two-tailed unpaired t-tests or Mann–Whitney tests were used; for multiple-group comparisons, one-way ANOVA with Tukey’s test or Kruskal–Wallis tests with Dunn’s test were used, according to data distribution. Exact *P* values are shown.

The accumulated TG reflected proportional expansion of the baseline pool. TGs with 56, 58, 54, and 60 total acyl carbons accounted for 38.4%, 31.0%, 15.4%, and 7.7% of the increase, respectively (Fig. 1d), while species with six, seven, or five double bonds accounted for 27.5%, 21.2%, and 17.6%, respectively (Fig. 1e). Each species’ contribution to newly accumulated TG tracked its baseline abundance without deviation from proportionality (slope ≈ 1.06, *P* = 0.12; Fig. 1f), and overall chain-length, unsaturation, and acyl-chain distributions were essentially unchanged (Extended Data Fig. 1f–h). Thus, OL2HG expands the TG pool in proportion to baseline abundance, rather than selectively remodeling its composition.

This TG-favoring lipotype was conserved across primary human cells. OL2HG increased total TG in endothelial cells and vascular smooth muscle cells, as in cardiomyocytes (*P* < 0.0001 in each; Fig. 1g). Differential expression of individual TG species and other lipids in cardiomyocytes, endothelial cells, vascular smooth muscle cells, and fibroblasts is provided in Supplementary Table 1.

In cardiomyocytes, both OL2HG and OD2HG also increased total DG, the immediate TG precursor (both *P* < 0.0001; Extended Data Fig. 1i), and 10 of 11 induced DG species shared their acyl chain composition with accumulated TGs (Extended Data Fig. 1j and Supplementary Table 1), consistent with progressive acylation to storage lipid. The similar DG increase but stronger TG induction with OL2HG compared with OD2HG suggests differential engagement of DGAT-mediated DG-to-TG acylation and/or TG clearance.

OL2HG also increased free oleate in all three cell types (*P* = 0.0037, 0.0195 and 0.0034; Fig. 1h), matching oleate as the dominant TG acyl constituent (∼40%; Extended Data Fig. 1h), without increasing the more lipotoxic free palmitate or stearate [except modestly so in vascular smooth muscle cells (Extended Data Fig. 1k,l)]. Cholesteryl esters showed a weaker and more cell-selective response, increasing at the class level in endothelial cells and fibroblasts (*P* < 0.0001 and 0.0448; Extended Data Fig. 1m), with some individual species also increased in cardiomyocytes and vascular smooth muscle cells (Supplementary Table 1). In addition, OL2HG markedly increased BODIPY 493/503- stained lipid droplets in cardiomyocytes (Fig. 1i,j) and Oil Red O-stained lipid droplets in vascular smooth muscle cells (Extended Data Fig. 1n,o). Thus, 2HG, most potently as L2HG, redirects fatty acyl groups into bulk TG and lipid droplets across several primary human cell types. We, therefore, focused on L2HG and next tested whether this TG-favoring lipotype reflects increased synthesis and/or slowed turnover.

### L2HG drives DGAT-mediated lipogenesis and slows TG turnover

To test DGAT dependence, we blocked terminal DG acylation with the DGAT1 inhibitor A-922500^25^ and the DGAT2 inhibitor PF-06424439^26^. DGAT inhibition abolished OL2HG-induced accumulation of BODIPY⁺ lipid droplets in oleate-loaded cardiomyocytes (Fig. 2a,b) and prevented the increase in total TG (OL2HG *vs* DMSO, *P* = 0.0001; with DGAT inhibition, *P* = 0.48; Extended Data Fig. 2a). Thus, L2HG-induced TG storage requires DGAT-mediated synthesis.

**Fig. 2:**
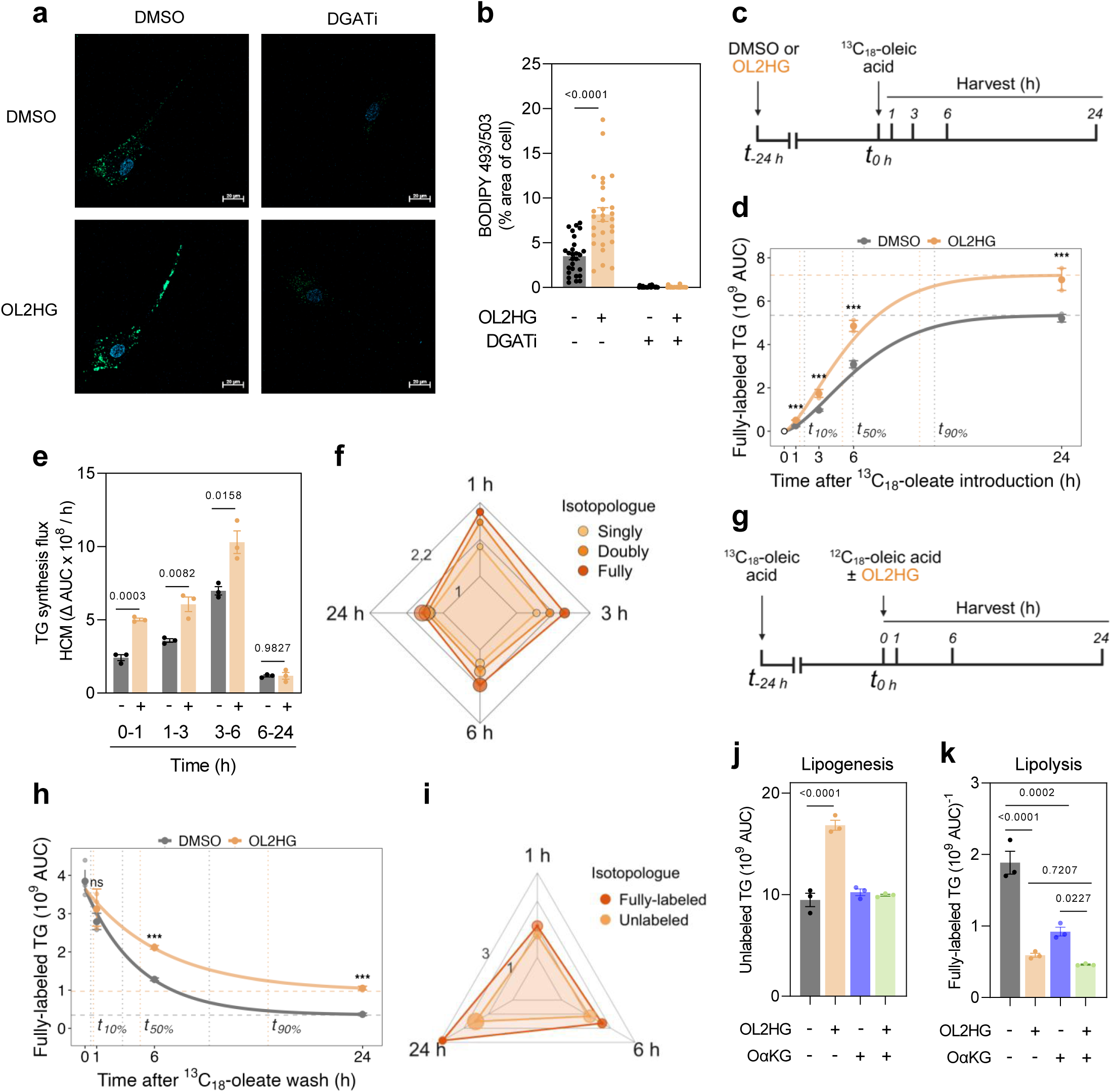
L2HG increases DGAT-dependent triglyceride synthesis and slows triglyceride turnover. **a,b,** Representative 63× BODIPY 493/503 images (**a**) and quantification (**b**) of oleate (50 µM)-loaded HCMs treated with DMSO or OL2HG (500 µM) ± combined DGAT1/2 inhibition (DGATi, A-922500 and PF-06424439, 10 µM each) for 24 h. Nuclei are stained with Hoechst. Scale bar, 20 µm; *n* = 27–28 cells from four independent samples per group. **c,** Pulse design: cells were pre-treated with DMSO or OL2HG (500 µM) for 24 h followed by [U-¹³C₁₈]-oleate (50 µM) labeling. **d,** Abundance of fully labeled triolein [TG(3 × ¹³C₁₈- 18:1)] during the pulse. Curves show through-origin Weibull fits; dashed lines indicate fitted plateau and time to 10%, 50%, and 90% of each condition’s plateau; *n* = 3 samples per condition per time point. **e,** Fully labeled triolein synthesis flux over the indicated intervals, calculated from **d**. **f,** Radar plot of OL2HG/DMSO labeling ratios for singly, doubly and fully labeled triolein isotopologues across the pulse in HCMs; node size indicates isotopologue abundance and opacity indicates significance. **g,** Pulse–chase design: cells were pulsed with [U-¹³C₁₈]-oleate (50 µM) for 24 h and chased with unlabeled oleate (50 µM) ± OL2HG (500 µM). **h,** Decay of pre-labeled fully labeled triolein during the chase in HCMs. Curve shows exponential-to-plateau fit; dashed lines indicate fitted plateau and time to 10%, 50% and 90% of each condition’s plateau; *n* = 3 samples per condition per time point, except t = 0 where *n* = 6. **i,** Radar plot of OL2HG/DMSO ratios for unlabeled and fully labeled triolein across the pulse–chase in HCMs; node size indicates isotopologue abundance and opacity indicates significance. **j,k,** Unlabeled (**j**) and fully labeled (**k**) triolein abundance at 24 h of chase in HCMs treated with OL2HG (500 µM) ± OαKG (500 µM). *n* = 3 per group. For flux curves (**d,h**), points show geometric mean ± geometric s.d.; *P* values were calculated by two-way ANOVA on log₁₀ AUC with Holm-adjusted OL2HG–DMSO contrasts within each time point. For all other panels, data are mean ± s.e.m. For two-group comparisons, two-tailed unpaired t-tests or Mann–Whitney tests were used; for multiple-group comparisons, one-way ANOVA with Tukey’s test or Kruskal–Wallis tests with Dunn’s test were used, according to data distribution. Exact *P* values are shown.

We quantified TG synthesis by pretreating cells with OL2HG or DMSO for 24 h, adding [U-¹³C₁₈]-oleate, and tracking labeled triolein [TG(3 × 18:1)] over 1–24 h (Fig. 2c). Because OL2HG expanded the unlabeled TG pool during pre-treatment, we used absolute labeled-isotopologue abundance rather than fractional enrichment. For fully labeled triolein [TG(3 × ^13^C18-18:1)], OL2HG increased the fitted labeling plateau 1.35-fold (nested-model F-test *P* = 0.023), accelerated filling (half-maximal labeling, 5.95 to 5.04 h; 90% labeling, 13.0 to 11.7 h), and increased labeled TG at every time point (1.3- to 2.0-fold; all *P* < 2 × 10⁻⁴; Fig. 2d). OL2HG also increased calculated synthesis flux over 0–1, 1–3, and 3–6 h (*P* = 0.0003, 0.0082, and 0.016), before convergence at 6–24 h as labeling approached isotopic steady state (Fig. 2e). The increase in maximal triolein labeling scaled across singly (^13^C18), doubly (^13^C36), and fully (^13^C54) labeled isotopologues (1.26-, 1.30-, and 1.35-fold; Extended Data Fig. 2b,c), and was resolved across isotopologues and time (Fig. 2f). OL2HG did not alter synthesis of fully labeled monoacylglycerol or diacylglycerol (Extended Data Fig. 2d,e), placing the effect at the terminal DGAT- catalyzed acylation step rather than the preceding acylation steps (although flux through the canonical GPAT–phosphatidic acid route was not traced directly). A similar increase in lipogenic flux occurred in endothelial cells, with a distinct temporal profile significant over the 0–1 h and 6–24 h windows (*P* = 0.0085 and 0.0005; Extended Data Fig. 2f,g).

Because TG synthesis and clearance can change independently, we performed the reciprocal pulse–chase experiment: cells were pre-labelled with [U-¹³C₁₈]-oleate for 24 h, then chased with unlabeled oleate ± OL2HG for 1–24 h (Fig. 2g). OL2HG slowed decay of pre- and fully-labeled triolein, extending its half-life from 3.2 to 4.8 h and increasing the fraction remaining at 24 h from 10% to 29% (decay F-test *P* = 1 × 10⁻¹¹; Fig. 2h). During the same chase, OL2HG increased newly synthesized unlabeled TG by ∼1.5-fold at 24 h (*P* = 8 × 10⁻⁵; Fig. 2i and Extended Data Fig. 2h), despite a transient, modest decrease at 1 h (0.80-fold, *P* = 0.009; Extended Data Fig. 2h). Thus, L2HG both slows TG turnover and increases sustained TG synthesis, with early time-dependent effects consistent with dynamic re-distribution of fatty acid partitioning. Notably, at 24 h, the surplus of newly synthesized unlabeled TG in OL2HG-treated cells (∼5.2 × 10⁹ AUC above DMSO; Extended Data Fig. 2h) exceeded the surplus of retained pre-labeled TG (∼6.9 × 10⁸ AUC above DMSO; Fig. 2h) by ∼sevenfold, indicating that increased synthesis is the dominant driver of TG expansion. To confirm that the retained triolein label reflected reduced clearance rather than recycling from other pre-labeled lipids, we repeated the chase under DGAT inhibition, and found that OL2HG-treated cells still retained more fully labeled triolein at 24 h (*P* = 0.0017; Extended Data Fig. 2i). Both slowed TG decay and increased unlabeled TG synthesis were reproduced in fibroblasts (Extended Data Fig. 2j,k).

αKG partially reversed the effects on the two components of the TG response. Specifically, octyl-esterified αKG (OαKG) reversed the OL2HG-induced increase in newly synthesized unlabeled triolein but potentiated the retention of pre-, fully-labeled triolein (Fig. 2j,k). Alone, OαKG did not increase oleate-to-TG flux but slowed TG turnover similarly to OL2HG (Fig. 2j,k). Thus, enhanced TG synthesis is specific to the reduced 2-hydroxy form, whereas slowed turnover is shared with αKG. We interpret the αKG effect cautiously, as it could reflect either opposition of an L2HG-specific target or competition at an αKG-dependent enzyme. Taken together, these data show that L2HG expands the TG pool by increasing DGAT-dependent synthesis and slowing TG turnover. Because DGAT draws on a diacylglycerol pool that is shared with Kennedy-pathway phospholipid synthesis, we next assessed whether L2HG-induced TG storage comes at the expense of phospholipids.

### L2HG diverts diacylglycerol flux away from phosphatidylethanolamine

The DGAT reaction consumes DG, the same intermediate used by the Kennedy pathway to synthesize phosphatidylethanolamine (PE) and phosphatidylcholine (PC). We, therefore, asked whether L2HG-driven triglyceride storage alters phospholipid abundance. In cardiomyocytes, OL2HG lowered total PE, measured by summing all detected PE species, as did OD2HG, although less strongly so (both *P* < 0.0001; Fig. 3a and Supplementary Table 1). Nearly half of the OL2HG-decreased PE species had acyl-chain compositions matching the increased TG species (19 of 39, 48.7%; Fig. 3b and Supplementary Table 2). This matched PE subset corresponds to ∼0.5 nmol acyl chains per well, comparable to the ∼0.36 nmol acyl chains newly incorporated into TG (Fig. 1c and Fig. 3a). Thus, although bulk pool sizes cannot establish species-level transfer, the composition and stoichiometry support diversion of a shared acyl-chain pool away from PE and toward TG. The PE decrease was reproduced in endothelial and vascular smooth muscle cells (*P* = 0.0027 and *P* < 0.0001; Fig. 3c). By contrast, total PC was unchanged by either enantiomer in cardiomyocytes (P > 0.09; Extended Data Fig. 3a). Accordingly, OL2HG, but not OD2HG, markedly reduced the PE/PC ratio (P < 0.0001 and P = 0.9542; Extended Data Fig. 3b), indicating selective PE depletion with preserved PC abundance.

**Fig. 3:**
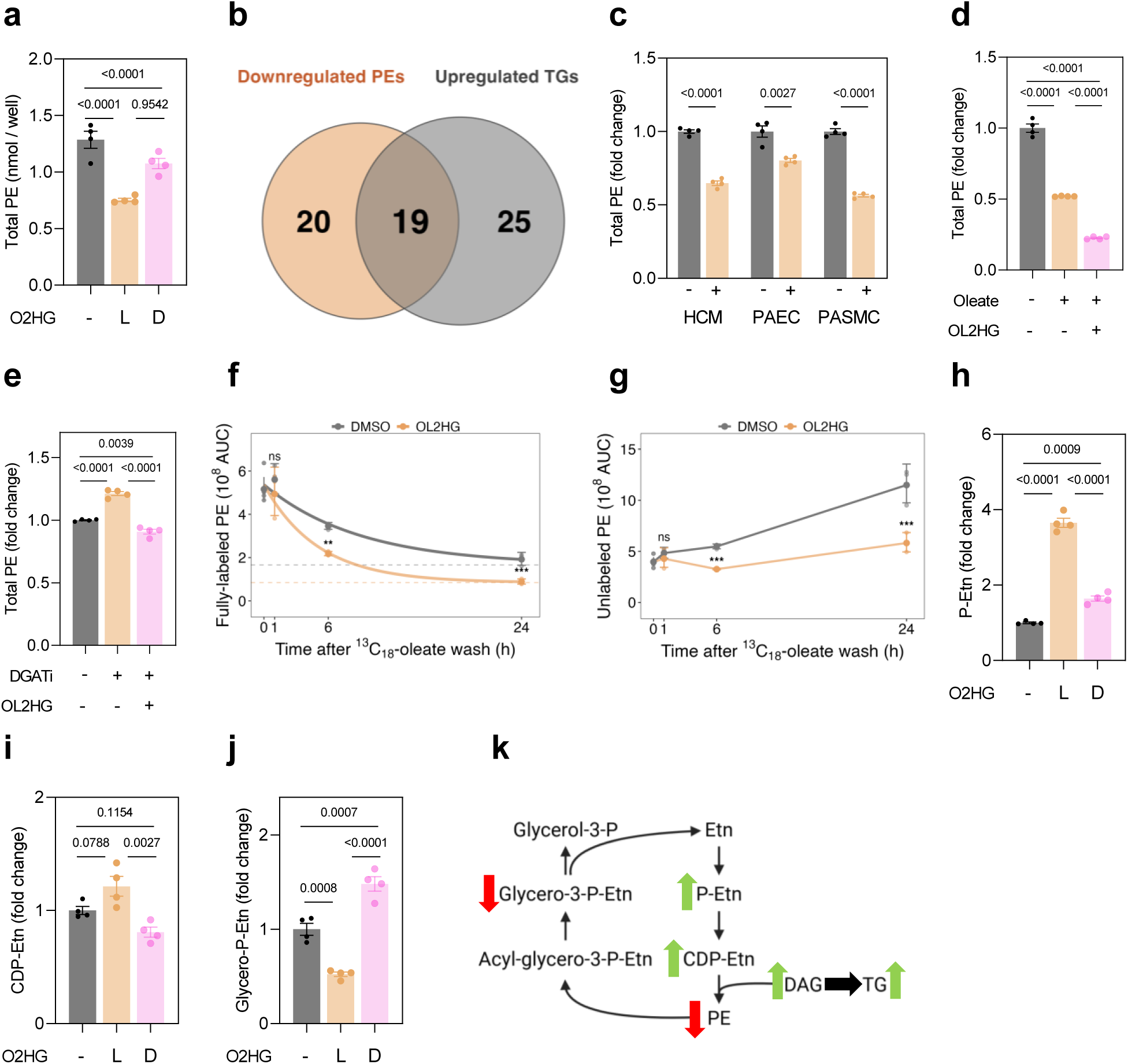
L2HG diverts DG flux away from PE. **a,** Total PE abundance in HCMs treated with DMSO, OL2HG (500 µM), or OD2HG (500 µM) for 24 h; *n* = 4 per group. **b,** Overlap between acyl-chain compositions of OL2HG-decreased PE species and OL2HG-increased TG species. Significance was defined as BH- adjusted *P* < 0.05 and ≥1.5-fold change. PE acyl pairs are unordered, *sn*-unassigned chain pairs; 19 of 39 decreased PE acyl pairs were also present in increased TG species. **c,** Total PE abundance in HCMs, PAECs, and PASMCs treated with DMSO or OL2HG (500 µM) for 24 h; *n* = 4 per group per cell type. **d,** Total PE abundance in PAECs treated with DMSO or oleate (50 µM) in the presence or absence of OL2HG (500 µM) for 24 h; *n* = 4 per group. **e,** Total PE abundance in PAECs treated with DMSO or combined DGAT1/2 inhibition (DGATi, A-922500 and PF-06424439, 10 µM each) in the presence or absence of OL2HG (500 µM) for 24 h; *n* = 4 per group. **f,g,** Clearance of fully labeled dioleoyl-PE [PE(2 × ^13^C18-18:1)] (**f**) and appearance of unlabeled dioleoyl-PE [PE(2 × ^12^C18-18:1)] (**g**) in HCMs pulsed with [U-¹³C₁₈]-oleate (50 µM) for 24 h and chased with DMSO or OL2HG (500 µM). *n* = 3 per condition per time point, except t = 0 where *n* = 6. Points show geometric mean ± geometric s.d.; *P* values were calculated by two-way ANOVA on log10 AUC with Holm-adjusted OL2HG–DMSO contrasts within each time point. **h–j,** Ethanolamine- branch Kennedy-pathway intermediates in HCMs treated with DMSO, OL2HG (500 µM), or OD2HG (500 µM) for 24 h: P-Etn (**h**), CDP-Etn (**i**), and glycero-P-Etn (**j**); *n* = 4 per group. **k,** Schematic of the L2HG-induced redirection of flux through the ethanolamine branch of the Kennedy pathway and TG synthesis. Unless stated otherwise, data are mean ± s.e.m. For two-group comparisons, two-tailed unpaired t-tests or Mann–Whitney tests were used; for multiple-group comparisons, one-way ANOVA with Tukey’s test or Kruskal–Wallis tests with Dunn’s test were used, according to data distribution. Exact *P* values are shown.

If OL2HG depletes PE by diverting DG into TG synthesis, DGAT inhibition should rescue PE. In endothelial cells, OL2HG potentiated oleate-induced PE loss (Fig. 3d), whereas DGAT inhibition increased basal PE and returned PE in OL2HG-treated cells toward baseline (Fig. 3e). This rescue was selective, as DGAT inhibition did not alter PC abundance (Extended Data Fig. 3c). Thus, OL2HG-induced PE depletion is coupled to DGAT-dependent TG synthesis, supporting diversion of DG toward storage at the expense of PE production.

We next used the [U-¹³C₁₈]-oleate pulse–chase design (Fig. 2g) to resolve PE synthesis and turnover. In cardiomyocytes, OL2HG decreased the retention of pre-, fully-labeled dioleoyl-PE [PE(2 x ^13^C18-18:1)] at 6 h and 24 h (0.63-fold, *P* = 0.0014; 0.46-fold, *P* = 1.6 × 10⁻⁵; Fig. 3f) and suppressed the accumulation of newly synthesized unlabeled dioleoyl-PE [PE(2 x ^12^C18-18:1)] at the same time points (0.59-fold, *P* = 6.4 × 10⁻⁴; 0.51-fold, *P* = 6.3 × 10⁻⁵; Fig. 3g). PC synthesis and clearance were unchanged (Extended Data Fig. 3d,e), reinforcing PE selectivity. Unlike TG, PE did not fit a simple rise-to-plateau kinetic model, consistent with continued remodeling during the chase rather than terminal storage. The same pattern was reproduced in fibroblasts, with reduced retention of pre-labeled PE at 6 h and 24 h (0.66-fold, *P* = 0.0024; 0.67-fold, *P* = 0.0036; Extended Data Fig. 3f), reduced accumulation of newly synthesized unlabeled PE (0.62-fold, *P* = 7.9 × 10⁻⁴; 0.60-fold, *P* = 5 × 10⁻⁴; Extended Data Fig. 3g), and unchanged PC flux (Extended Data Fig. 3h,i). At 24 h, the deficit in newly synthesized PE exceeded the deficit in retained pre-labeled PE by ∼fivefold in both cardiomyocytes (∼5.7 × 10⁸ *vs* ∼1.0 × 10⁸ AUC; Fig. 3f,g) and fibroblasts (∼4.9 × 10⁸ *vs* ∼1.0 × 10⁸ AUC; Extended Data Fig. 3f,g), indicating that impaired PE synthesis, more than accelerated turnover, primarily drives PE depletion.

We localized this constraint by quantifying ethanolamine-branch Kennedy-pathway intermediates. OL2HG increased phospho-ethanolamine (P-Etn) ∼3.8-fold in cardiomyocytes (*P* < 0.0001; Fig. 3h) and produced a trend toward increased CDP-Etn, the immediate substrate for PE synthesis (*P* = 0.079; Fig. 3i), while reducing the PE-derived catabolite glycero-P-Etn by approximately half (*P* = 0.0008; Fig. 3j). Each change was greater in magnitude with L2HG than D2HG. Accumulation of P-Etn and CDP-Etn, together with reduced glycero-P-Etn, was reproduced largely in fibroblasts and endothelial cells (Extended Data Fig. 3j–o). In endothelial cells, OL2HG preserved labeling of ¹³C₂-P- Etn from [¹³C₂]-ethanolamine but reduced labeling of downstream ¹³C₂-CDP-Etn and ¹³C₂- glycero-P-Etn (Extended Data Fig. 3p–r), consistent with impaired ethanolamine flux through the PE branch of the Kennedy pathway. Thus, L2HG couples DGAT-dependent TG storage to selective PE depletion and Kennedy pathway constraint (Fig. 3k).

### L2HG decreases lipid peroxidation, suppresses fatty acid oxidation, and decreases lipotoxic acylcarnitines

L2HG-induced fatty acid sequestration away from PE and into TG also reduced lipid peroxidation. OL2HG lowered the oxidized-to-reduced ratio of BODIPY-C11 in cardiomyocytes (*P* = 0.036), and DGAT inhibition abolished this effect (*P* = 0.86; Extended Data Fig. 4a,b). We next asked whether the same storage shift limits fatty acid availability for mitochondrial β-oxidation.

OL2HG also limited fatty acid oxidation, reducing ¹⁴C-palmitate oxidation in cardiomyocytes (*P* = 0.028; Fig. 4a) and in endothelial cells (*P* = 0.014; Fig. 4b). Long-chain acylcarnitines, mitochondrially lipotoxic intermediates of incomplete β-oxidation, decreased in a DGAT-dependent manner in cardiomyocytes, most prominently oleoylcarnitine [CAR(18:1)] and palmitoylcarnitine [CAR(16:0)], followed by stearoylcarnitine [CAR(18:0)] (Fig. 4c).

**Fig. 4:**
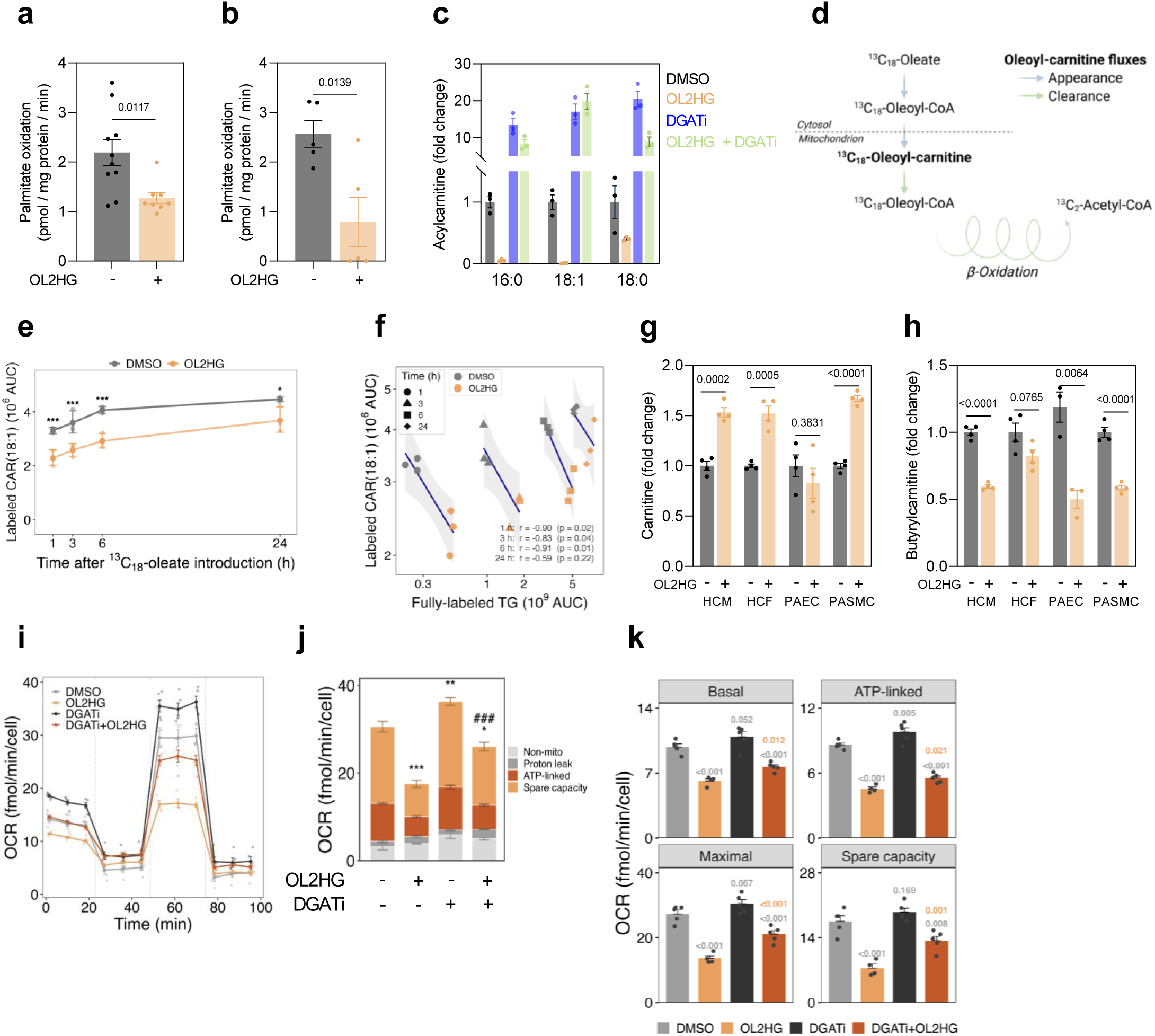
L2HG suppresses fatty acid oxidation and lowers lipotoxic acylcarnitines. **a,b,** ¹⁴C-palmitate oxidation in HCMs (**a**) and PAECs (**b**) treated with DMSO or OL2HG (500 µM) for 24 h. **c,** Long-chain acylcarnitines [CAR(16:0), CAR(18:1) and CAR(18:0)] in HCMs treated with DMSO, OL2HG (500 µM), DGATi (A-922500 and PF-06424439, 10 µM each) or OL2HG + DGATi for 24 h; *n* = 3 per group. **d,** Schematic of [U-¹³C₁₈]- oleate entry into oleoylcarnitine and mitochondrial β-oxidation. **e,** ^13^C18-oleoylcarnitine abundance in HCMs pre-treated with DMSO or OL2HG (500 µM) for 24 h followed by [U-¹³C₁₈]-oleate labeling (50 µM); *n* = 3 per condition per time point. Significance was tested by two-way ANOVA with Holm-adjusted OL2HG–DMSO contrasts within each time point. **f,** Per-well correlation between ^13^C18-oleoylcarnitine and fully labeled triolein [TG(3 × ¹³C₁₈-18:1)]. *n* = 6 per time point. **g,h,** Free carnitine (**g**) and butyrylcarnitine (**h**) abundance in HCMs, HCFs, PAECs and PASMCs treated with DMSO or OL2HG (500 µM) for 24 h; *n* = 4 per group per cell type. **i–k,** Seahorse OCR during the mitochondrial stress test in HCMs treated with DMSO, OL2HG (500 µM), DGATi (A-922500 and PF-06424439, 10 µM each) or OL2HG + DGATi for 24 h; *n* = 4–5 per group. Significance was tested by one-way ANOVA using emmeans. Unless otherwise stated, data are mean ± s.e.m.. For two-group comparisons, two-tailed unpaired t-tests or Mann–Whitney tests were used; for multiple-group comparisons, one-way ANOVA with Tukey’s test or Kruskal–Wallis tests with Dunn’s test were used, according to data distribution. Exact *P* values are shown.

Next, using the design shown in Fig. 2c, we traced [U-¹³C₁₈]-oleate into oleoylcarnitine, a marker of oleate commitment to mitochondrial import and/or oxidation (Fig. 4d). OL2HG lowered labeled oleoylcarnitine at every measured time point in cardiomyocytes (0.69- to 0.83-fold, *P* ≤ 0.018; Fig. 4e), and labeled oleoylcarnitine decreased as labeled TG increased across individual wells (Pearson r = −0.83 to −0.91 from 1–6 h; Fig. 4f). This suppression was reproduced in endothelial cells and fibroblasts with cell type-specific kinetics (Extended Data Fig. 4c,d). The storage–oxidation balance was also time-dependent: in the pulse–chase design (Fig. 2g), OL2HG increased newly synthesized unlabeled oleoylcarnitine at 1 h in cardiomyocytes and fibroblasts before later suppression (Extended Data Fig. 4e,f), internally consistent with the early decrease and later increase in unlabeled TG synthesis (Extended Data Fig. 2h). Thus, L2HG shifts fatty acid partitioning dynamically, with the sustained 24 h state favoring TG storage and reduced oleoylcarnitine formation. Moreover, OL2HG increased free carnitine in cardiomyocytes, fibroblasts, and vascular smooth muscle cells (∼1.3- to 1.7-fold, *P* ≤ 0.0005), but not in endothelial cells (Fig. 4g), while reducing butyrylcarnitine and acetylcarnitine (Fig. 4h and Extended Data Fig. 4g). Butyrylcarnitine suppression persisted after OL2HG washout despite intracellular 2HG falling from ∼20-fold to ∼5-fold above DMSO within 3 h (Extended Data Fig. 4h,i), indicating a durable shift rather than simple proportionality to prevailing 2HG abundance. Etomoxir partially phenocopied this signature by lowering propionylcarnitine and butyrylcarnitine, with OL2HG further decreasing butyrylcarnitine (Extended Data Fig. 4j). OL2HG also decreased extracellular free carnitine and butyrylcarnitine in cardiomyocyte and fibroblast medium (Extended Data Fig. 4k,l). The hallmark fatty acid metabolism pathway was enriched in OL2HG-treated cells despite decreased oxidative flux, and this enrichment was reversed by DGAT inhibition (NES 1.98 to 1.23; Extended Data Fig. 4m). Taken together, decreased long- and short-chain acylcarnitines with increased intracellular free carnitine indicates reduced fatty acid entry into, or throughput through, β-oxidation, consistent with decreased ¹⁴C-palmitate oxidation and slower TG clearance.

L2HG-induced lipid partitioning also altered mitochondrial energetics. Seahorse respirometry showed that OL2HG suppressed basal, ATP-linked, maximal, and spare respiration in cardiomyocytes (all *P* < 10⁻⁴; Fig. 4i–k), while increasing glycolysis (∼1.8- fold basal extracellular acidification, *P* < 10⁻⁶; Extended Data Fig. 4l,m). DGAT inhibition partially restored respiration, most clearly maximal and spare capacity (Fig. 4k), but did not reverse the glycolytic increase (Extended Data Fig. 4n,o), indicating that TG storage contributes to loss of respiratory reserve whereas glycolytic rewiring is storage independent.

### L2HG-induced phospho- and redox-proteotype are partially dependent on TG accumulation

We next used tandem mass tag (TMT) phosphoproteomics to define OL2HG signaling and its dependence on DGAT. Across 4,229 quantified phosphosites, OL2HG altered 109 sites (78 increased, 31 decreased; *q* < 0.05 and |FC| ≥ 1.5; Extended Data Fig. 5a and Supplementary Table 3). Kinase activity inference showed suppression of proline-directed CDK and ERK signaling and activation of basophilic AGC kinase programs, including PKC, ROCK, and PKA (Extended Data Fig. 5b and Supplementary Table 3). Most OL2HG-responsive phosphosites were DGAT-independent (99 of 109), including suppressed cell-cycle–associated sites on NPM1-Thr199, NCL-Thr121, TOP2A-Ser1393, RB1-Ser807, and PTTG1-Ser165 (Extended Data Fig. 5c and Supplementary Table 3). These findings suggest that 2HG may influence oncogenic signaling not only through chromatin regulation, but also through broad remodeling of kinase-dependent phosphorylation networks. Interestingly, OL2HG increased phosphorylation of BCKDHA at Ser347, revealing a potential mechanism that may contribute to our previously established link between branched-chain ketoacids and L2HG^27^. Notably, seven sites were reversed by DGAT inhibition, defining a small storage-coupled module that included the glycolytic enzyme GAPDH-Thr211 and the choline/ethanolamine transporter SLC44A2- Thr691, linking TG storage to signaling nodes potentially relevant to the glycolytic rewiring and ethanolamine/PE Kennedy pathway remodeling that we report herein. Phospho-ubiquitin Ser65, a PINK1-linked marker of mitochondrial stress and mitophagy initiation^28^, emerged only with DGAT inhibition and increased ∼12-fold with combined OL2HG and DGAT inhibition *vs.* DMSO, suggesting that OL2HG-driven TG expansion protects against mitochondrial stress when lipid storage capacity is intact.

The cysteine-redox proteotype induced by OL2HG showed greater dependence on TG storage. Using a TMT-multiplexed thiol-tagging assay, we quantified the free-thiol fractional occupancy of 5,881 high-confidence cysteines (quantified by ≥ 2 independent spectra) and identified 108 OL2HG-responsive sites (87 more oxidized, 21 more reduced; |Δ| ≥ 10 percentage points, *P* < 0.01, excluding cysteines with non-physiological occupancy, *i.e.*, % reduced > 110% in either condition; Extended Data Fig. 5d and Supplementary Table 3). Nearly half were DGAT-dependent, including sites on mitochondrial and antioxidant proteins such as LONP1 Cys909 and GPX1 Cys78/Cys156. By contrast, FASN showed DGAT-independent redox changes at Cys779 and Cys1759, placing the lipogenic enzyme itself under OL2HG-linked redox control. Thus, OL2HG signaling at the phosphoproteome level is largely independent of TG storage, whereas DGAT-dependent storage couples to a distinct mitochondrial and antioxidant redox arm.

Taken together, these data identify L2HG as a suppressor of fatty acid oxidation that limits accumulation of long-chain acylcarnitines, modulates relevant proteins in the phospho- and redox-proteomes, and limits mediators of lipotoxic stress by routing fatty acids into storage. Because coordinated lipid-partitioning programs are often transcriptionally regulated, we next asked whether L2HG acts through canonical transcriptional regulators of lipid storage.

### L2HG’s induced regulators are dispensable for TG storage

OL2HG induced a strong pseudohypoxic transcriptional program, but this program was not required for the lipotype. Hallmark enrichment placed hypoxia at the top of both the OL2HG-regulated transcriptome and proteome in cardiomyocytes (Extended Data Fig. 6a–d and Supplementary Table 4). HIF-1α was the most activated inferred transcription factor, whereas E2F factors were most repressed (Extended Data Fig. 6e and Supplementary Table 4), and immunoblotting confirmed HIF-1α stabilization by OL2HG (Extended Data Fig. 6f). Importantly, however, *HIF1A* silencing did not prevent OL2HG-induced TG accumulation, oleoylcarnitine depletion, or PE loss (Extended Data Fig. 6g–j). Conversely, hypoxia and *HIF1A* transcriptional programs remained enriched when OL2HG-induced TG storage was blocked by DGAT inhibition (Extended Data Fig. 6k). Thus, L2HG- induced lipid partitioning and the pseudohypoxic transcriptional response are mutually independent.

Canonical lipid-storage regulators were also dispensable. OL2HG increased perilipin 2 (PLIN2), a lipid droplet coat protein, and very-low density lipoprotein receptor (VLDLR), a receptor for triglyceride-rich lipoproteins, in the proteome (Extended Data Fig. 7a), with PLIN2 induction reversing upon DGAT inhibition and VLDLR remaining storage-independent (Extended Data Fig. 7b). PLIN2 induction was specific relative to PLIN3 and PLIN5 (Extended Data Fig. 7c,d), and occurred at the transcript level in cardiomyocytes and in endothelial cells (Extended Data Fig. 7e–f); yet, *PLIN2* silencing did not prevent the OL2HG-induced TG accumulation (Extended Data Fig. 7g–i). OL2HG also increased VLDL uptake (Extended Data Fig. 7j–k), but not LDL uptake (Extended Data Fig. 7l,m). Removing extracellular lipoproteins or serum lowered absolute TG abundance but preserved the OL2HG-induced fold increase in TG and PLIN2 induction (Extended Data Fig. 7n–q), indicating that extracellular lipid supply contributes to basal TG abundance but is not required for the L2HG-mediated storage response.

SREBP1 was similarly dispensable. OL2HG stabilized SREBP1, but not SREBP2, without inducing *SREBF1* mRNA in cardiomyocytes, and this response was reproduced in endothelial cells (Extended Data Fig. 8a–f). *SREBF1* silencing did not prevent OL2HG-induced TG accumulation or PLIN2 induction (Extended Data Fig. 8g–i). Thus, L2HG builds TG stores independently of pseudohypoxic transcription, PLIN2, VLDLR, and SREBP1.

### L2HG induces hypertriglyceridemia, redistributes the TG pool along acyl chain length, and limits ischemic lipotoxicity

To test whether L2HG promotes lipid storage in vivo, we profiled plasma from *L2hgdh^−/−^*mice, which accumulate L2HG because they cannot degrade it. *L2hgdh^−/−^* mice showed ∼7- to 11-fold higher circulating 2HG and 1.4- to 2-fold higher total plasma TG in both sexes (Extended Data Fig. 9a,b), with 19 TG species increased by 1.5- to 1.8-fold (Fig. 5a and Supplementary Table 5). As in cells, TG expansion was proportional rather than compositional: the increase scaled with baseline abundance, spanned chain length, and unsaturation, and preserved overall acyl composition (Fig. 5b–d and Extended Data Fig. 9c–e).

**Fig. 5:**
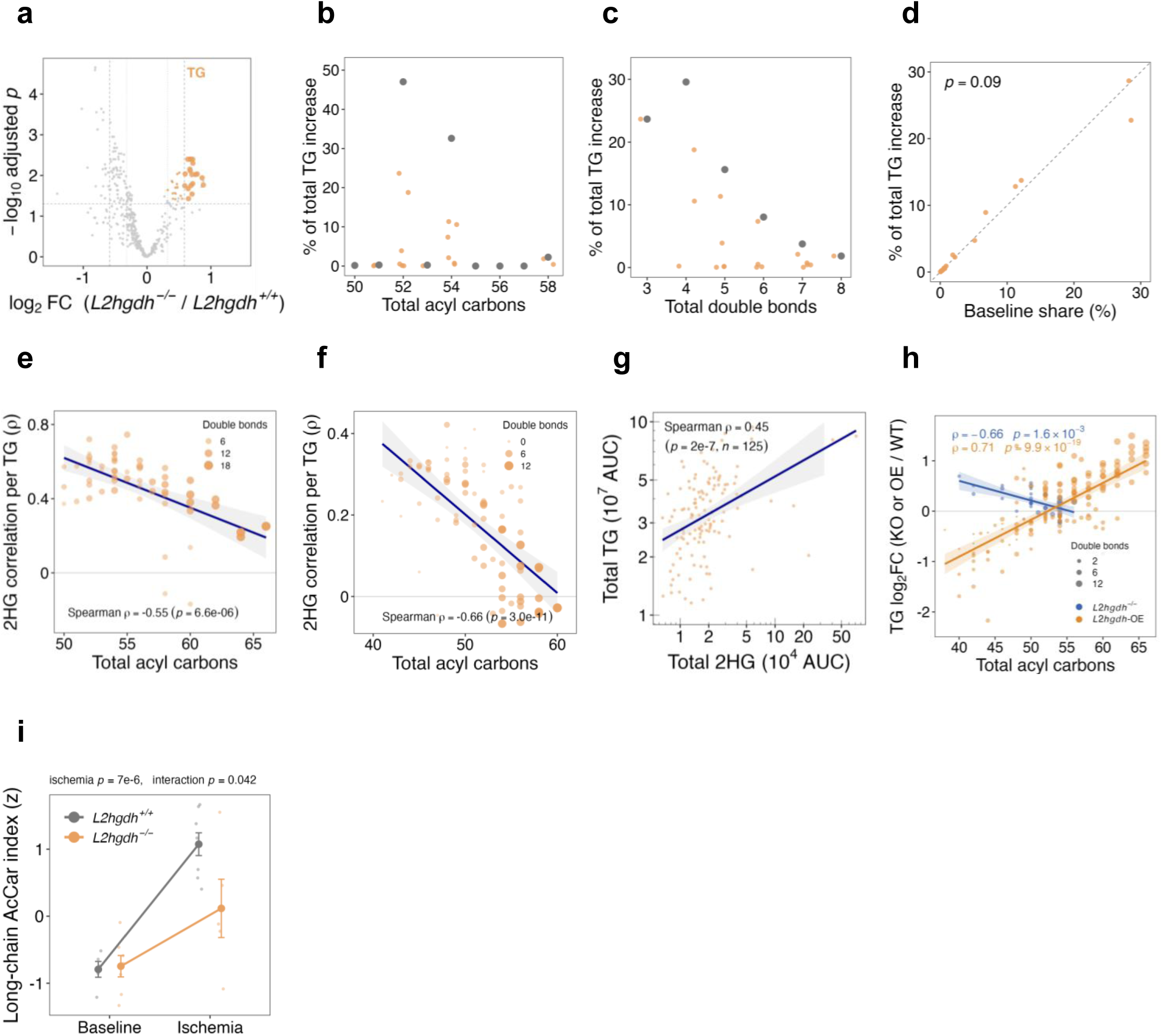
*L2hgdh–/–* mice exhibit hypertriglyceridemia and attenuated long-chain acylcarnitine accumulation during myocardial ischemia. **a**, Volcano plot of plasma lipidomics from *L2hgdh^−/−^* mice (*n* = 10 equally split across sex) *vs. L2hgdh^+/+^* mice (*n* = 12 equally split across sex). TG species are orange; other lipids are grey. Significance: |FC| ≥ 1.5 and BH-adjusted *P* < 0.05. **b,c,** Contribution of each plasma TG species to the *L2hgdh^−/−^*-associated TG increase by total acyl-carbon number (**b**) and total double bonds (**c**); orange, individual TG species; grey, summed bin. **d,** Per- species contribution to newly accumulated TG versus baseline pool share among the 19 increased TG species. Dashed line, y = x; slope = 0.93, *P* = 0.09 by two-sided t-test of the fitted regression slope against 1, indicating proportional expansion. **e,f,** Per-species Spearman correlation between plasma 2HG and TG abundance plotted against total acyl carbon number in mice [**e**; 59 TG species across *n* = 22 mice (*L2hgdh^−/−^*, *n* = 10 equally split across sex; *L2hgdh^+/+^*, *n* = 12 equally split across sex)] and humans (**f**; 79 TG species across *n* = 714 individuals). **g,** Spearman correlation between plasma 2HG and TG (≤ 50 total acyl carbons) in the Columbia Bariatric and Diabetes human weight-loss cohort; *n* = 125 individuals (see Data availability). **h,** Per-species TG fold change plotted against total acyl chain carbon number in *L2hgdh^−/−^* (blue, *n* = 8 *L2hgdh^−/−^* and *n* = 12 *L2hgdh^+/+^*) and *L2hgdh*-overexpressing (orange, *n* = 5 per genotype) hearts *vs.* their respective wild-type control animals. Solid lines show linear regression with 95% confidence intervals. **i**, Long-chain acylcarnitine index, calculated as the mean *z*-score of the detected long chain acylcarnitines, in Langendorff-perfused hearts from male *L2hgdh^−/−^* and *L2hgdh^+/+^* mice at baseline and after ischemia; *n* = 5–8 hearts per group. Significance was tested by two-way ANOVA. Unless otherwise stated, data are mean ± s.e.m. For two-group comparisons, two-tailed unpaired t-tests or Mann–Whitney tests were used; for multiple-group comparisons, one-way ANOVA with Tukey’s test or Kruskal–Wallis tests with Dunn’s test were used, according to data distribution. Exact *P* values are shown.

The circulating 2HG–TG relationship was strongest for shorter-chain TGs. Across individual TG species, the correlation between plasma 2HG and TG abundance weakened with increasing acyl-chain length and degree of unsaturation in mice (Spearman ρ = −0.55, *P* = 6.6 × 10^-6^; Fig. 5e) and humans (ρ = −0.66, *P* = 3.0 × 10^-11^; Fig. 5f). Circulating 2HG also correlated positively with the shorter-chain TG pool, defined as TG species with ≤50 total acyl carbons, across three independent human cohorts (ρ = 0.45, 0.30, and 0.29; *P* ≤ 2 × 10⁻⁴; Fig. 5g and Extended Data Fig. 9f,g), supporting a conserved association between 2HG and TG storage across mice and humans.

Although baseline total TG abundance in both hearts and livers were unchanged in *L2hgdh^−/−^* and *L2hgdh* overexpressing mice (which have depleted systemic L2HG^19^) (Extended Data Fig. 9h,i), L2HG selectively reshaped the cardiac TG pool by acyl chain length. This effect was absent in liver and reversed with L2HG abundance: *L2hgdh^−/−^* hearts shifted proportionally toward shorter TG species (ρ = −0.66, *P* = 1.6 × 10^-3^), whereas *L2hgdh*-overexpressing hearts shifted proportionally toward longer, more polyunsaturated species (ρ = +0.71, *P* = 9.9 × 10^-19^), suggesting that, under metabolic steady state, L2HG regulates which acyl chains are stored as cardiac TG without changing total TG abundance or significantly altering individual species (Fig. 5h and Extended Data Fig. 9j). Because lipid peroxidation increases with double-bond content^29^, which generally rises with acyl-chain length, the shorter, more saturated TG species that correlate with circulating and cardiac 2HG are intrinsically less susceptible to peroxidation.

Finally, because L2HG protects mice against myocardial ischemia^30^, we asked whether elevated L2HG limits acute ischemic lipotoxicity. In *ex vivo* Langendorff-perfused hearts, ischemia increased long-chain acylcarnitines, a signature of incomplete fatty-acid oxidation^31^, and this response was blunted in *L2hgdh^−/−^* hearts (interaction *P* = 0.042; Fig. 5i). Attenuation was significant for palmitoylcarnitine and hydroxylinoleoylcarnitine, with a concordant trend for linoleoylcarnitine (interaction *P* = 0.045, 0.026 and 0.121; Extended Data Fig. 9k–m). Thus, elevated L2HG limits the generation of ischemia-induced lipotoxic intermediates, extending the metabolite’s regulation of fatty acid metabolism to the ischemic heart.

Conversely, re-analysis of a published *L2hgdh*-overexpression RNA-seq dataset^19^ showed that fatty acid and TG metabolism programs decreased coordinately in kidney and liver (NES −1.3 to −2.7; Extended Data Fig. 9n), a response opposite in direction to, and internally consistent with, our observation in OL2HG-treated cells (Extended Data Fig.4k).

## Discussion

Here, we identify L2HG and D2HG as metabolites that determine fatty acid fate. Rather than changing fatty acid abundance alone, 2HG changes where fatty acids are routed, biasing them toward triglyceride storage and away from membrane phospholipid synthesis and mitochondrial oxidation. This partitioning converges on diacylglycerol, the shared substrate for DGAT-dependent triglyceride synthesis and Kennedy-pathway phospholipid synthesis. By increasing DGAT-dependent triglyceride synthesis while slowing triglyceride turnover, 2HG redirects this shared intermediate into storage at the expense of phosphatidylethanolamine, which is depleted while phosphatidylcholine is spared (Fig. 6). A metabolite that gates this branch point toward storage, selectively depleting one membrane phospholipid class while preserving another, has, to our knowledge, not been described previously.

**Fig. 6:**
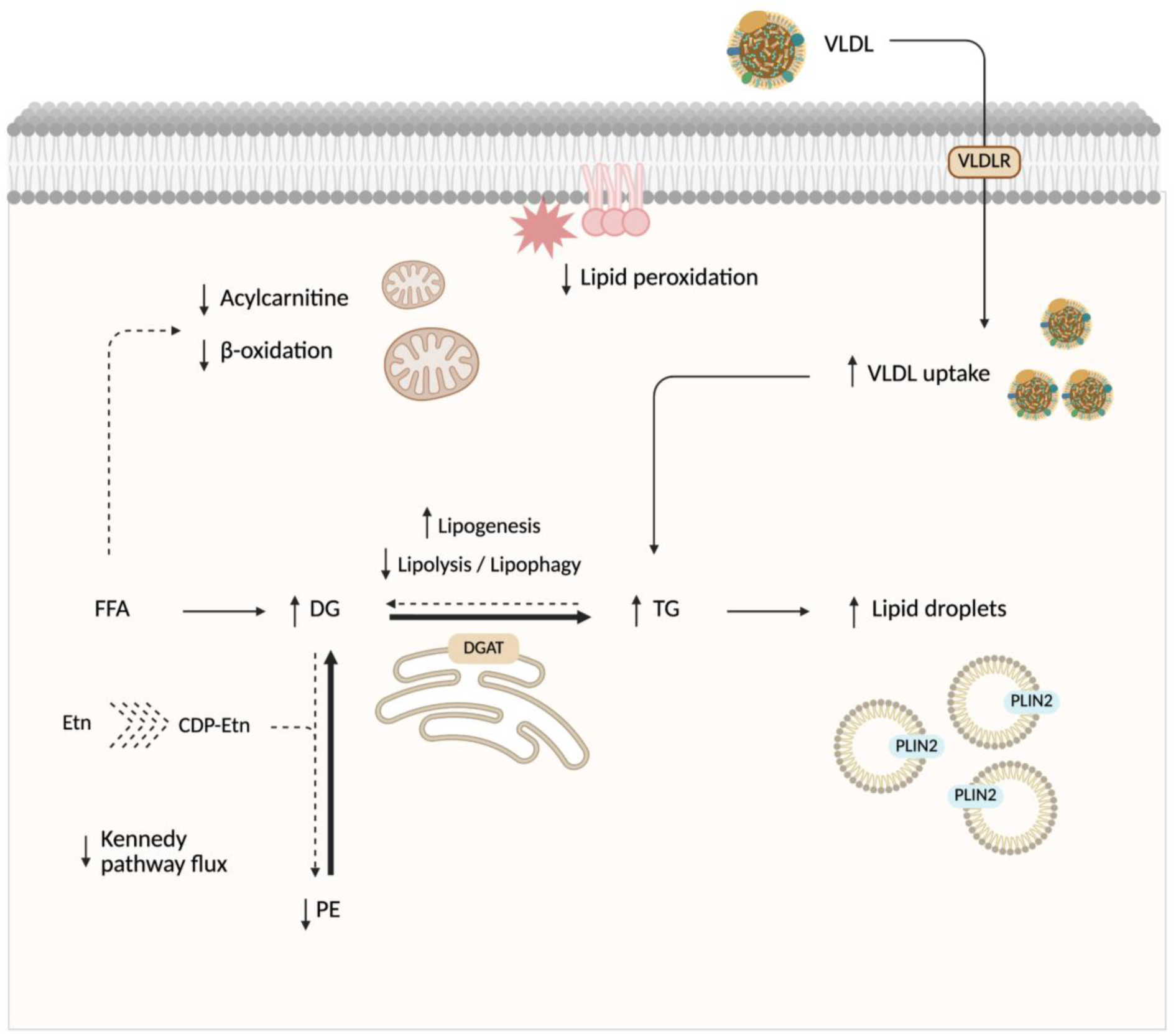
Model of L2HG-induced lipid remodeling. Graphical summary of the lipid-partitioning program induced by L2HG, and, to a lesser extent, D2HG. Bold arrows denote increased flux and dashed arrows denote decreased flux. L2HG increases intracellular TG and lipid droplets by coordinately increasing DGAT-dependent TG synthesis (lipogenesis) and decreasing TG clearance (lipolysis and/or lipophagy). At the DG branch point, this shift diverts flux away from the ethanolamine branch of the Kennedy pathway, decreasing PE synthesis and increasing PE turnover, thereby lowering cellular PE. In parallel, L2HG suppresses fatty acid β-oxidation, lowers long-chain acylcarnitines, and reduces lipid peroxidation. L2HG also increases VLDL uptake and induces PLIN2 expression, although both are dispensable for the TG accumulation phenotype. Taken together, these findings support a model in which 2HG limits lipotoxicity by redirecting fatty-acid fate away from membrane phospholipid synthesis and mitochondrial oxidation and toward neutral lipid storage.

The notion that that fatty acid fate is not fixed, but partitioned among storage, oxidation, and membrane synthesis through reciprocally coupled pathways is a recognized metabolic pattern. Triglyceride synthesis protects cells by sequestering excess fatty acids that would otherwise become lipotoxic^3^, yet the stored pool remains metabolically available when needed. Lipophagy releases lipid-droplet fatty acids for β-oxidation, and blocking this process increases triglyceride content^32^. Adipose triglyceride lipase generates lipid ligands that activate the PPARα/PGC-1α oxidative program, such that impaired lipolysis suppresses fatty-acid-oxidation gene expression and promotes triglyceride accumulation^33^. This storage–oxidation exchange is spatially organized at lipid droplet–mitochondria interfaces, where mitochondrial dynamics route droplet-derived fatty acids between oxidation and re-esterification^34^, and where DGAT1-dependent lipid droplet biogenesis buffers autophagy-derived fatty acids to protect mitochondria from lipotoxic overload^5^. DGAT1-mediated re-esterification also counterbalances lipolysis in a futile cycle that limits fatty acid-induced endoplasmic-reticulum stress^35^. Storage is coupled to membrane synthesis, as well, because localized phosphatidylcholine production is required for lipid droplet expansion, making phospholipid supply a determinant of storage capacity^36^. Our findings place 2HG upstream of this tripartite system, biasing fatty acids toward DGAT-dependent triglyceride storage and away from both β-oxidation and the phosphatidylethanolamine arm of membrane synthesis. Thus, 2HG acts not on a single lipid fate, but as a set-point regulator of the storage–oxidation–membrane balance.

Our work adds 2HG to the set of endogenous metabolites that govern fatty acid partitioning, but through a distinct logic. Several metabolites are recognized as lipid fate signals. Malonyl-CoA allosterically inhibits carnitine palmitoyltransferase 1 to limit mitochondrial fatty acid entry and favor esterification^37^. Citrate activates acetyl-CoA carboxylase to promote de novo lipogenesis^38^, while acetyl-CoA supplies lipogenic carbon and, through ATP-citrate lyase, links nutrient state to histone acetylation and lipogenic gene expression^39^. Other metabolites act through signaling pathways to affect lipid metabolism: succinate inhibits adipocyte lipolysis through SUCNR1^40,41^, cholesterol and oxysterols control SCAP/INSIG-dependent activation of SREBP^42^, and *S*-adenosylmethionine regulates the PE-to-PC ratio as the methyl donor for PEMT such that excess methylation can deplete PE^43^. 2HG differs from these paradigms. It is not a lipogenic substrate, a known allosteric regulator of a lipid enzyme, or a receptor ligand for lipolytic control. Instead, it is an α-ketoglutarate-derived signaling metabolite that biases flux at the DGAT–Kennedy branch point, a node not previously recognized as directly gated by a central carbon metabolite, coupling DGAT-dependent triglyceride storage to selective PE depletion. In addition, this effect is at least partly specific to the 2-hydroxy form, as equimolar α-ketoglutarate alone did not induce lipogenic flux. Overall, 2HG extends the catalogue of lipid-fate metabolites as a non-canonical, (onco)metabolite-class regulator of fatty acid partitioning.

Our findings are likely to have broad relevance because 2HG accumulates in multiple physiological and pathological settings characterized by lipid remodeling. In clear-cell renal cell carcinoma, L2HG accumulates through chromosome-14q loss of *L2HGDH*^20^, while HIF2α–PLIN2 signaling drives triglyceride storage in support of tumor survival^44^. In IDH1/IDH2-mutant gliomas and leukemias, D2HG reaches low-millimolar concentrations^11^ and remodels phospholipid metabolism, including phosphatidylethanolamine^45,46^. Hypoxia likewise induces L2HG production^16,17^ in parallel with HIF-dependent lipid droplet formation^47^; L2HG regulates CD8⁺ T-cell fate^21^, a process influenced by DGAT1-mediated triglyceride storage^48^; and the ischemic myocardium accumulates L2HG^30^ while also suppressing fatty acid oxidation and lipid remodeling^49^. Most directly, mitochondrial dysfunction elevates D2HG and redirects fatty acids into lipid droplets to mediate whitening of brown adipocytes^50^. Taken together, these observations support a broader model in which 2HG acts as a conserved metabolic regulator of fatty acid partitioning across cancer, hypoxia, immunity, and ischemia settings.

Several questions remain. First, the molecular target linking 2HG to the DGAT–Kennedy branch point remains unknown, including whether it is an α-ketoglutarate-dependent enzyme or another protein engaged by covalent^23^ or non-covalent interaction^24^. Second, although the induced transcriptional programs and lipid storage regulators we tested were dispensable, we did not globally block transcription or chromatin-modifying dioxygenases and, therefore, cannot exclude contributions from untested transcriptional or epigenetic mechanisms. Third, our pulse–chase experiments measured net TG turnover but did not distinguish lipolysis from lipophagy^51^, nor did they trace direct transfer of acyl chains from PE to TG. Because [U-¹³C₁₈]-oleate labels both pools simultaneously, definitive species-level transfer would require selective PE labeling followed by chase into TG. Fourth, our flux analyses focused on oleate; other fatty acids may be partitioned differently by 2HG depending on chain length and saturation. Fifth, most experiments focused on sustained 24 h 2HG exposure, when TG storage was maximal and most reproducible. The opposite early response at 1 h (*i.e.*, reduced labeled TG with increased labeled oleoyl-carnitine) indicates that 2HG-dependent fatty-acid partitioning is dynamic rather than fixed, and likely depends on exposure duration, substrate availability, and metabolic state. Finally, how the cell-autonomous TG retention observed here relates to circulating hypertriglyceridemia *in vivo*, through altered hepatic secretion, peripheral clearance or both, remains to be defined.

Overall, our findings argue for a broader view of metabolic flexibility. Flexibility is usually framed as the capacity to switch between fuels, most classically glucose and fatty acids^7^. Yet, each fuel is itself partitioned among competing fates: glucose can be oxidized, stored as glycogen, or diverted into serine and one-carbon metabolism; glutamine can support anaplerosis, nucleotide, protein, and glutathione synthesis, or reductive carboxylation toward lipogenesis; and fatty acids are allocated among oxidation, storage, and membrane synthesis. By identifying 2HG as a diffusible signal that redirects fatty acid flux toward DGAT-dependent triglyceride storage and away from phospholipid synthesis and mitochondrial oxidation, our work establishes fate-level fuel partitioning as a regulated dimension of metabolic flexibility.

## Data availability

Differential-expression results underlying all omics analyses are provided in Supplementary Tables 1–5. The *L2hgdh*-overexpression RNA-seq re-analyzed in Extended Data Fig. 9n is from Ref.^19^ and is available from the Gene Expression Omnibus under accession GSE326291. The human plasma 2-hydroxyglutarate and triglyceride measurements analyzed in Fig. 5f,g and Extended Data Fig. 9f,g are from a publicly available plasma metabolomics dataset reported in Ref.^52^ from three weight loss intervention cohorts (The Weight Loss Maintenance, WLM^53^, NCT00054925; The Studies of Targeted Risk Reduction Interventions through Defined Exercise in individuals with Pre-Diabetes, STRRIDE-PD^54,55^, NCT00962962; and The Columbia Bariatric and Diabetes, CBD^56–60^, NCT01516320, NCT02287285, NCT02929212, NCT00571220) and is openly available at Zenodo (https://doi.org/10.5281/zenodo.4767969). The mass spectrometry lipidomics data and the RNA-sequencing data generated in this study have been deposited at [repository] and [repository], respectively, under accession [to be assigned] and [to be assigned], respectively. The mass spectrometry proteomics data have been deposited to the ProteomeXchange Consortium via the PRIDE^61^ partner repository with the dataset identifier PXD081894. Source data are provided with this paper.

## Acknowledgements

The authors thank Rose Cannistraro for expert technical support. This work was supported in part by US National Institutes of Health grants R01 HL155107 and R01 HL166137, American Heart Association MERIT grant 1185447, EU HorizonHealth grant 10157619 (RePO4EU), and a generous gift from the Ruggeri family to JL.

## Methods

### Chemicals and reagents

LC–MS grade water (cat. no. LC365), methanol (cat. no. 34966), and acetonitrile (cat. no. LC015), were purchased from Honeywell. LC–MS-grade chloroform (cat. no. AC364320010), isopropanol (cat. no. A461), and iodoacetamide (cat. no. 122270050) were purchased from Fisher Scientific. *d8*-valine (cat. no. DLM-311), ^13^C5-glutamine (cat. no. CLM-1822), ^13^C2-ethanolamine (cat. no. CLM-274-PK), and ^13^C18-oleic acid (cat. no. 490431) were purchased from Cambridge Isotope Laboratories. ^14^C1-palmitic acid (cat. no. NEC075H050UC) was purchased from Revvity. LC–MS vials (cat. no. 5182-0716) with 250 µL inserts (cat. no. 5183-2085) were purchased from Agilent. Butylated hydroxytoluene (cat. no. B1378), ammonium carbonate (cat. no. 379999), medronic acid (cat. no. 64255), ammonium formate (cat. no. 70221), unmodified L2HG (cat. no. 90790), dithiothreitol (cat. no. D0632), sodium dodecyl sulfate (SDS; cat. no. 151213), and oleic acid (cat. no O1008) were purchased from Sigma. EquiSPLASH™ (cat. no. 330731) and Carnitine SPLASH™ (cat. no. 330379) were purchased from Avanti Polar Lipids. Octyl-L2HG (OL2HG, cat. no. 16367), octyl-D2HG (OD2HG, cat. no. 16366), octyl-α-ketoglutarate (OαKG, cat. no. 11970), carnitine (cat. no. 21489), BSA–palmitate complex (cat. no. 29558), etomoxir (cat. no. 11969), PF-06424439 (cat. no. 17680), and A-922500 (cat. no. 10012708) were purchased from Cayman Chemical.

### Cell culture and treatments

Primary human cardiac myocytes (HCM) isolated from the ventricles of an adult heart (cat. no. C-12810, PromoCell) were cultured in complete myocyte growth medium (cat. no. C-22070, PromoCell). Primary human pulmonary artery endothelial cells (PAEC; cat. no. CC-2530, Lonza) were cultured in endothelial basal medium (cat. no. CC-3156, Lonza) with supplements (cat. no. CC-4176, Lonza). Primary human ventricular cardiac fibroblasts (HCF; cat. no. CC-2904, Lonza) were cultured in fibroblast growth medium (cat. no. CC-3131, Lonza) with supplements (cat. no. CC-45525, Lonza). Primary human pulmonary arterial smooth muscle cells (PASMC; cat. no. CC-2581, Lonza) were cultured in SmBM® Basal Medium (cat. no. CC-3181, Lonza) and SmGM®-2 SingleQuots® supplements (cat. no. CC-4149, Lonza). Passages 4–8 were used for all primary cell types. The cells used in this study were obtained from commercial sources and were marketed by the suppliers as originating from non-diseased donors.

All cells were maintained at 37 °C in an incubator with 95% humidity and 5% CO2. Medium was refreshed every other day for cell growth. For treatments, cells were refreshed with new media supplemented with the appropriate treatment for 24 h at the following concentrations, unless otherwise indicated in the main text or Figure legends: OL2HG (500 µM), OD2HG (500 µM), octyl acetate (0.25–1 mM), unmodified L2HG (0.5–2 mM), A-922500 (10 µM), PF-06424439 (10 µM). For controls, an equal volume of sterile dimethyl sulfoxide (DMSO, cat. no. D2650, Sigma) was used, except where otherwise indicated where ultrapure DNase/RNase-free distilled water (cat. no. 10977015, Thermo Fisher) was used as an additional/alternative control.

### Targeted metabolomics, non-targeted lipidomics, and stable-isotope–based fluxomics

All solvents outlined below are LC–MS-grade.

#### Sample preparation

For metabolomic analyses, cells were plated in 6-well plates at a concentration of 2.5 x 10^5^ cells per well with the exception of HCM, which were plated at 1.5 x 10^5^ cells per well. For lipidomic analyses, cells were plated in 12-well plates at a concentration of 1.25 x 10^5^ cells per well with the exception of HCM, which were plated at 7.5 x 10^4^ cells per well. Cells were treated as described in the main text and figure legends, washed twice with LC– MS-grade water, flash-frozen by placing the plate on liquid N2, and stored at -80 °C until LC–MS analysis as described below.

Metabolites were extracted from frozen cells, tissues, or plasma/medium (50 µL) by adding 200 μL of extraction buffer [50/30/20 (v/v/v) methanol/acetonitrile/water containing 1 μM *d8*-valine and1 μM ^13^C5-glutamine] to each well of the plate on ice. After ∼10 min, lysates were scraped, collected, and centrifuged (17,000 x *g*, 10 min, 4 °C). The supernatant (180 μL) was concentrated to dryness at 45 °C using a refrigerated CentriVap vacuum concentrator equipped with a cold trap (Labconco). Dried extracts were kept at -80 °C. Immediately before LC–MS analysis, extracts were reconstituted in 50 μL of water, centrifuged (17,000 x *g*, 10 min, 4 °C), and 35 μL of the supernatant transferred to glass LC–MS vials for analysis. For each experimental batch, an additional 10 μL of the supernatant from each sample were pooled, centrifuged (17,000 x *g*, 10 min, 4 °C), and transferred to glass LC–MS vials for quality control purposes.

To extract lipids from frozen cells, tissues, or plasma (50 µL), 200 μL of 1/1 (v/v) methanol/water were added to each well of the plate on ice. After ∼10 min, lysates were scraped and collected. To each lysate, 50 μL of butylated hydroxytoluene (1 mg/mL in methanol), 5 μL of an internal standard mixture [1/1/38 (v/v/v) EquiSPLASH™/Carnitine SPLASH™ /methanol), and 200 μL of chloroform were added. The mixtures were mixed vigorously for 10 s, centrifuged (17,000 x *g*, 10 min, 4 °C), and 180 μL of the bottom chloroform layer were collected and concentrated to dryness under a gentle stream of N2 gas. Dried extracts were kept at -80 °C. Immediately before LC–MS analysis, extracts were reconstituted in 100 μL of 1/1 (v/v) chloroform/methanol, centrifuged (17,000 x *g*, 10 min, 4 °C), and 80 μL of the supernatant were transferred to amber glass LC–MS vials for positive and negative acquisition, as described below. For each experimental batch, an additional 10 μL of the supernatant from each sample were pooled, centrifuged (17,000 x *g*, 10 min, 4 °C), and transferred to glass LC–MS vials for quality control purposes.

#### LC–MS data acquisition and analysis

LC–MS analyses were performed using a Vanquish ultra-high-performance liquid chromatography (UHPLC) system connected to a Q Exactive Orbitrap mass spectrometer with a heated electrospray ionization (HESI)-II source (Thermo Fisher Scientific). Samples were stored in the autosampler at 5 °C prior to injection. For metabolomics, the following chromatographic parameters were used: solvent A, 20 mM ammonium carbonate in water containing 5 μM medronic acid, pH corrected to 9.2 with ammonium hydroxide; solvent B, acetonitrile; injection volume, 2 μL; oven temperature, 25 °C; flow rate, 0.1 mL/min. Compounds were separated using a SeQuant zwitterionic polymer-based hydrophilic interaction liquid chromatography (ZIC-pHILIC) column (5 μm, 150 × 2.1 mm; Merck) with a guard column and a linear gradient of solvent B as follows: 0 min, 80%; 20 min, 20%; 20.5 min, 8%; 24 min, 8%; 24.5 min; 80%; 35 min, 80%. MS scan parameters were as follows: scan type, full MS; scan range, 60-900 Th; fragmentation, none; resolution, 70,000; microscans, 1; lock masses, off; automatic gain control (AGC) target, 1 x 10^6^; maximum injection time, 80 ms. The MS was operated in polarity switching mode. HESI source parameters were as follows: sheath gas flow rate, 40, auxiliary gas flow rate, 15; sweep gas flow rate, 1; spray voltage, 1 kV; capillary temperature, 320 °C; S-lens radio frequency level, 50; auxiliary gas heater temperature, 350 °C. Targeted peak identification was performed using the Thermo TraceFinder General LC software (Thermo Fisher Scientific), using an in-house library of ∼150 metabolites with known retention times, as assessed and optimized *a priori* with authentic standards using the same method parameters.

For lipidomics, the following chromatographic parameters were used: solvent A, 10 mM ammonium formate in 60/40 (v/v) acetonitrile/water; solvent B, 10 mM ammonium formate in 90/10 (v/v) isopropanol/acetonitrile; injection volume, 5 μL; oven temperature, 55 °C; flow rate, 0.3 mL/min. Compounds were separated using an ACQUITY UPLC charged surface hybrid (CSH) C18 column (1.7 μm, 100 × 2.1 mm; Waters) with a guard column and a linear gradient of solvent B as follows: 0 min, 0%; 6 min, 40%; 30 min, 100%; 34 min, 100%; 36 min; 0%; 40 min, 0%. Data were acquired using data-dependent MS2 acquisition. MS1 scan parameters were as follows: scan type, full MS; scan range, 200-1200 Th; fragmentation, none; resolution, 70,000; microscans, 1; lock masses, off; AGC target, 1 x 10^6^; maximum injection time, 250 ms. MS2 scan parameters were as follows: scan type, AIF; scan range, 200-1200 Th; fragmentation, collision energy, 30; resolution, 17,500; microscans, 1; lock masses, off; AGC target, 1 x 10^5^; maximum injection time, 120 ms; loop count, 5; TopN, 5; isolation window, 1.0 m/z; first fixed mass, 50 m/z. Data dependent (dd) settings were as follows: maximum AGC target, 5 x 10^3^; intensity threshold, 4.2 x 10^4^; apex trigger, off; charge exclusion, off; peptide match, off; exclude isotopes, off; dynamic exclusion, 20 s. Positive and negative acquisition modes were used separately. HESI source parameters were as follows: sheath gas flow rate, 40 for negative mode and 50 for positive mode, auxiliary gas flow rate, 15 for negative mode and 7 for positive mode; sweep gas flow rate, 1 for negative mode and 5 for positive mode; spray voltage, 3 kV; capillary temperature, 320 °C for negative mode and 300 °C for positive mode; S-lens RF level, 70; auxiliary gas heater temperature, 350 °C for negative mode and 300 °C for positive mode. Non-targeted lipidomic analysis was performed using LipidMatch (v5.62; Innovative Omics) with blank filtering. Only the features with the highest confident identifications (*i.e.*, confirmed by MS2) were used for subsequent statistical analyses. For stable isotope flux experiments with [U-^13^C18]-oleate, peaks of interest were integrated manually using the software Freestyle (Thermo Fisher Scientific).

For both metabolomics and lipidomics, unless otherwise stated, peak intensities for each feature were normalized to the total metabolite pool (sum of integrated peak intensities across all identified targeted features) or total lipid pool (sum of integrated peak intensities across all high-confidence, MS2-confirmed lipid features), respectively. Cell viability and morphology were monitored visually at harvest in all experiments; none of the treatments produced detectable changes in adherence or morphology across the primary cell types used. In experiments where treatments were anticipated to affect viability, protein-content normalization was applied *a priori*.

For lipidomics, analyses used only the highest-confidence identifications [MS2-confirmed lipid features (LipidMatch v5.62 rank-1)]. Sum-normalized peak areas were log2-transformed, and differential abundance between treatments was tested with limma (v3.64.1) moderated t-statistics (lmFit followed by empirical-Bayes moderation with the trend and robust options) under a design contrasting the treatment groups, with Benjamini–Hochberg (BH) control of the false-discovery rate across the pooled positive- and negative-ionization features; species were called significant at *q* < 0.05 and |log2 fold-change| ≥ log2(1.5). The contribution of each species to the treatment-induced change was expressed as its share of the summed change in abundance (100 × ΔAUCi / ΣΔAUC). Analyses were performed in R v4.5.0 with ggplot2 (v4.0.2).

### Lipogenesis, lipolysis, and acylcarnitine flux experiments

Stable-isotope tracing was used to quantify flux through TG synthesis and fatty acid handling using two complementary experimental designs. In both approaches, oleic acid uniformly labeled with heavy carbon ([U-^13^C18]-oleate) was supplied at 50 µM, and lipid species were quantified by LC–MS based on isotopologue composition with emphasis on triolein containing zero [TG(3 × ^12^C18-18:1)], one [TG(2 × ^12^C18-18:1; 1 × ^13^C18-18:1)], two [TG(2 × ^13^C18-18:1; 2 × ^12^C18-18:1)], or three [TG(3 × ^13^C18-18:1)] labeled oleate chains, as well as the corresponding unlabeled and labeled diolein, monoolein, dioleoyl-PE, dioleoyl-PC, and oleoylcarnitine.

The first approach was designed to assess labeling/appearance kinetics. Cells were pre-treated with vehicle (DMSO) or OL2HG (500 µM) for 24 h, after which complete medium was refreshed containing [U-^13^C18]-oleate (50 µM) (Fig. 2c). Cells were harvested at variable time points between 1 and 24 h following tracer addition. Lipogenic flux from extracellular oleate into intracellular TG stores and other lipid fates was inferred from the time-dependent appearance of different isotopologues of triolein, diolein, monoolein, dioleoyl-PE, dioleoyl-PC, and oleoylcarnitine.

In the second approach, a classic pulse–chase design was implemented. Cells were first pulsed with complete medium containing [U-^13^C18]-oleate (50 µM) for 24 h to pre-label intracellular lipid pools. Media were then replaced with fresh medium containing equimolar unlabeled oleate ([U-^12^C18]-oleate) together with either DMSO or OL2HG (500 µM), and cells were harvested across a 0–24 h chase (Fig. 2g). Turnover/clearance of pre-labeled lipid pools was quantified using LC–MS as described above from the clearance kinetics of labeled species, whereas concurrent synthesis and/or recycling under chase conditions was assessed from the appearance of newly synthesized unlabeled species over time.

For each condition, the absolute peak area of every ^13^C-labeled isotopologue was modeled over time by non-linear least-squares regression on log10-transformed signal, with the functional form (Weibull rise through the origin for label appearance; single-exponential approach-to-plateau for label clearance) selected by the Akaike information criterion; the effect of condition on the fitted trajectory was tested by nested-model F-tests. At individual time points, isotopologue abundances were compared by two-way ANOVA (condition × time) on log10(peak area) with Holm-adjusted estimated-marginal-means contrasts (emmeans v2.0.2), and relationships between labeled pools (Fig 4f) were quantified by Pearson correlation on log10 values. Analyses were performed in R v4.5.0.

### BODIPY staining

HCMs were seeded in 8-well chamber slides (cat. no. C8SB-1.5H, Cellvis) at 10,000 cells per well and allowed to adhere overnight in a humidified incubator (37 °C, 95% humidity, 5% CO₂). Cells were treated as described in the text or figure legends, washed once with 300 µL serum-free myocyte growth medium, and stained with either BODIPY 493/503 (4,4-Difluoro-1,3,5,7,8-Pentamethyl-4-Bora-3a,4a-Diaza-s-Indacene; 2 µg/mL; cat. no. D3922, Thermo Fisher Scientific) to quantify lipid droplets or with BODIPY 581/591 C11 (2 µg/mL; cat. no. D3861, Thermo Fisher Scientific) to quantify lipid peroxidation. Staining was performed in serum-free myocyte growth medium for 30 min in a dark humidified incubator (37 °C, 95% humidity, 5% CO₂). Following staining, cells were washed twice with Hanks’ Balanced Salt Solution (HBSS; cat. no. 14025076, Thermo Fisher Scientific) and refreshed with 300 µL HBSS containing 1 µg/mL Hoechst (cat. no. 62249, Invitrogen) for 5 min. Cells were washed twice with 300 µL HBSS and images were acquired on a Zeiss LSM 800 confocal microscope using identical settings across all treatments.

Ratiometric image acquisition and fluorescence intensity quantification of BODIPY C11 were performed using ZEN software (blue edition; Carl Zeiss Microscopy, Jena, Germany) on an LSM 800 confocal microscope. BODIPY 493/503 stained images were segmented per cell and the percentage of cell area occupied by droplets were quantified using an in-house script in ImageJ. Statistics were produced in R v4.5.0 with ggplot2 (v4.0.2).

### Oil Red O staining

Cells were stained using Oil Red O staining kit (cat. no. SKU: LL-0052, Lifeline Cell Technology). PASMCs were seeded in 6-well plates at 200,000 cells per well, allowed to adhere overnight, and treated with DMSO, OL2HG (500 µM), or OD2HG (500 µM) for 24 h. Following treatment, cells were washed twice with DPBS, fixed, dehydrated, and stained according to the manufacturer’s instructions. Color bright field images were acquired at 40**×** magnification using an Agilent BioTek Cytation Imaging Reader (cat. no. CYT5FV, Agilent Technologies). Images were quantified in Python 3.10 using scikit-image 0.25.2, NumPy 2.2.6, Pillow 9.4.0, and SciPy 1.15.3. For each field, the burned-in scale bar was excluded by masking the lower 7% of rows and rightmost 30% of columns.

Per-channel background intensity was estimated from the histogram mode of the unstained substrate and used to convert each image to optical density, OD = −log10(I/I₀), thereby correcting for field-to-field differences in white balance and illumination. Punctate lipid droplets were separated from diffuse nonspecific staining by white top-hat morphological filtering with a disk structuring element of radius 12 pixels. Droplet-positive pixels were defined using a single global threshold, set as the mean plus three standard deviations of the top-hat signal pooled across all vehicle-treated fields and applied uniformly to all images. Objects smaller than 16 pixels were removed. Oil Red O droplet content was expressed as the percentage of field area occupied by droplet-positive pixels.

### Lipoprotein uptake

Purified human very low-density lipoprotein (VLDL) and low-density lipoprotein (LDL) labeled with 1,1’-dioctadecyl-3,3,3’,3’- tetramethylindocarbocyanine perchlorate (Dil-VLDL and Dil-LDL, cat. no. 770130-9 and 770230-9, respectively, Kalen Biomedical) were diluted aseptically in serum-free HCM medium immediately prior to use. HCMs were seeded at 10,000 cells per well in 8-chamber slides and cultured overnight in complete HCM medium.

Cells were treated for 24 h as described in the text or figure legends. Following treatment, cells were incubated with either DiI-VLDL (50 µg/mL, 2 h) or Dil-LDL (10 µg/mL, 3 h) in serum-free HCM medium in a dark humidified incubator (37 °C, 95% humidity, 5% CO₂). Cells were washed twice with warm serum-free medium and counterstained with Hoechst (1 µg/mL) for 5 min in HBSS.

Fluorescence images were acquired using a Zeiss LSM 800 confocal microscope (excitation 549 nm, emission 565 nm) with identical acquisition settings for all intra-batch samples. Quantification was performed using ImageJ. Data are presented as mean fluorescence intensity (MFI) per field normalized to cell.

### Palmitate oxidation

Palmitate oxidation was quantified by measuring the production of radiolabeled CO2 from [^14^C]-palmitate, as adapted from Ref^62^. HCMs and PAECs were seeded in 24-well plates (25,000 and 50,000 cells per well, respectively) and allowed to adhere overnight. Cells were treated with either vehicle control (DMSO) or 500 µM OL2HG for 24 h before assessment of palmitate oxidation. For palmitate oxidation, [^14^C]-palmitate (cat. no. NEC075H050UC) was added to 5 mM cold palmitate (cat. no. 29558) (exact reagent volumes were back-calculated for each experiment based on the number of wells used). This palmitate mixture was incubated at 37 °C for 60 min with gentle shaking to ensure complete complex formation and diluted into serum-free MCDB 131 medium (cat. no. CM134-050) to final medium concentrations of 200 µM cold palmitate and 0.4 µCi mL⁻¹ [^14^C]-palmitate. Carnitine (cat. no. 21489) was added to this radiolabeled BSA–palmitate mixture to a final concentration of 1 mM. Cells were washed twice with DPBS and incubated with the radiolabeled palmitate-containing medium for 3 h at 37 °C in an incubator with 95% humidity and 5% CO2, with plates sealed using parafilm. Following incubation, 400 µL of medium was transferred to 1.5 mL tubes containing 200 µL of 1 M perchloric acid, with filter paper (cat. no. WHA1004270) pre-soaked with 20 µL of 1 N NaOH mounted in the tube cap. After incubation at room temperature for 1 h while shaking at 200 r.p.m, filters containing trapped ^14^CO2 were transferred to scintillation vials filled with 4 mL of Ecoscint H (cat. no. LS-275), mixed vigorously for 10 s, and radioactivity was measured as disintegrations per minute (DPM) using a liquid scintillation counter (LS 6500, Beckman Coulter). Non-incubated medium samples were processed in parallel to determine input radioactivity for calculation of specific activity.

All procedures involving radioactive materials were performed in accordance with institutional radiation safety guidelines and approved protocols at Brigham and Women’s Hospital and Harvard Medical School (Boston, MA, USA).

### Mitochondrial respirometry and glycolytic flux (Seahorse)

Oxygen consumption rate (OCR) and extracellular acidification rate (ECAR) were measured with a Seahorse XFe24 extracellular flux analyzer (Agilent Technologies). HCMs were seeded in Seahorse XF24 cell culture microplates (cat. no. 100777-004) at 25,000 cells per well and treated for 24 h with vehicle (DMSO), OL2HG (500 µM), DGAT1/2 inhibitors (A-922500 and PF-06424439, 10 µM each), or DGAT inhibitors + OL2HG. On the day of the assay, growth medium was replaced with Seahorse XF assay medium supplemented with 10 mM D-Glucose, 2 mM L-Glutamine, and 1 mM pyruvate, and cells were equilibrated for 1 h at 37 °C in a non-CO2 incubator. A mitochondrial stress test was performed by sequential injection of oligomycin (1 µM), the uncoupler BAM15 (1 µM), and rotenone plus antimycin A (0.5 µM each). Basal respiration, ATP-linked respiration, proton leak, maximal respiration and spare respiratory capacity were derived from the OCR trace, and basal glycolysis from the ECAR; values were normalized to cell number.

Respiratory and glycolytic parameters were compared across the four conditions by two-way ANOVA (OL2HG × DGAT inhibitor) with Holm-adjusted estimated-marginal-means contrasts (emmeans v2.0.2); DGAT-dependence of each parameter was evaluated from the OL2HG × DGAT-inhibitor interaction term and quantified as the fraction of the OL2HG effect reversed by DGAT inhibition. Analyses were performed in R v4.5.0.

### RNA sequencing

HCMs and PAECs were plated in 6-well plates at 150,000 cells per well or 250,000 cells per well, respectively, and allowed to adhere overnight. Cells were then treated with vehicle (DMSO) or OL2HG (500 µM) for 24 h (*n* = 4 replicates per condition). Following treatment, cells were washed twice with LC–MS-grade water and total RNA was isolated using the miRNeasy kit (cat. no. 217604, Qiagen) according to the manufacturer’s instructions. RNA concentration and purity were assessed by UV spectrophotometry using a NanoDrop (cat. no. 3400518, Thermo Scientific). RNA samples were shipped to Innomics Inc. (Cambridge, MA, USA) for library preparation, sequencing, and primary data processing. Processed expression data were returned as counts or transcript-per-million (TPM) matrices and were subsequently aggregated at the gene level.

Sequencing reads were quantified to gene-level counts (Salmon for the HCM libraries and RSEM for the PAEC libraries), and differential expression between OL2HG and vehicle was tested with DESeq2 (v1.48.2; negative-binomial generalized linear model with Wald test) using vehicle as the reference; genes were called significant at BH-adjusted *P* < 0.05 with apeglm-shrunken (v1.30.0) |log2 fold-change| ≥ log2(2). Pathway enrichment was assessed by pre-ranked gene-set enrichment analysis (fgsea v1.34.2) over the MSigDB Hallmark, C2 (canonical pathways) and C5 (Gene Ontology) collections (msigdbr v26.1.0), ranking genes by the DESeq2 Wald statistic. Transcription factor activities were inferred with decoupleR^63^ (v2.14.0; univariate linear model) over the CollecTRI regulons^64^ (retrieved *via* OmnipathR v3.16.2 from OmniPath on 6 June 2026; 1,178 TFs and 42,601 signed interactions), using the DESeq2 Wald statistic as input. Analyses were performed in R v4.5.0 / Bioconductor 3.21.

### LC–MS-based proteomics

#### Global proteomics and phosphoproteomics sample preparation

HCMs were plated in 60-mm dishes at 450,000 cells per dish and treated for 24 h with DMSO or OL2HG (500 µM) ± DGATi (A-922500 and PF-06424439, 10 µM each). Cells were washed twice with LC–MS-grade water and immediately lysed by syringe lysis and tip probe sonication in 8 M urea in 200 mM EPPS buffer (pH 8.5, corrected with ammonium hydroxide) supplemented with 1 × Halt protease and phosphatase inhibitors (cat. no. 78440). Protein concentration was determined by bicinchoninic acid (BCA) assay. Samples were reduced with dithiothreitol (10 mM, 30 min, room temperature) and alkylated with iodoacetamide (20 mM, 1 h, room temperature, protected from light). Samples were then standardized to 200 µg total protein, precipitated by chloroform– methanol fractionation, and digested with Trypsin/Lys-C (25:1, overnight, 37 °C; cat. no. PR-V5073, Promega). Peptides were labeled with 16-plex TMTpro reagents (cat. no. A44521, Thermo Fisher Scientific) across 12 channels (100 µg peptide per channel at a 1:2.5 peptide:label ratio, 1 h, room temperature). Labeling efficiency was confirmed to be >98% by LC–MS/MS on a pooled aliquot before final pooling of the 12 channels using different volumes to account for signal ratios. The pooled sample was then quenched with hydroxylamine (0.3%, v/v, final concentration), acidified to 1% (v/v) TFA, and desalted by reversed-phase solid-phase extraction using a 100 mg Sep-Pak cartridge (cat. no. WAT023590, Waters).

Phosphopeptides were enriched from the pooled TMT-labeled peptides by Fe-NTA immobilized-metal-affinity chromatography using the single-enrichment “mini-phos” strategy as described previously^65^, with the High-Select Fe-NTA Phosphopeptide Enrichment Kit (cat. no. A32992, Thermo Fisher Scientific). The unbound flow-through and first wash were pooled and fractionated as described below for the global proteome. Enriched phosphopeptides were desalted by reversed-phase solid-phase extraction using a 100 mg Sep-Pak cartridge and analyzed on the Orbitrap platform as described below.

#### Cysteine-redox proteomics sample preparation

HCMs were plated in 10-cm dishes and grown to ∼80% confluency. Cells were treated for 24 h with DMSO or OL2HG (500 µM) ± DGATi (A-922500 and PF-06424439, 10 µM each) (*n* = 3 per condition). To preserve the native cysteine redox state, the lysis buffer (8 M urea, 1 mM EDTA, 200 mM EPPS pH 8.5, with 1 × Halt protease and phosphatase inhibitors) and the LC–MS-grade water used for washing the cells were sparged with N2 for 20–30 min. Cells were washed twice with N2-sparged water and syringe lysed (without tip-probe sonication, to avoid peroxide-driven artefactual oxidation) on ice. Each lysate was split into two identical 75 µg aliquots. In the total-thiol arm, reversibly oxidized thiols and disulfides were reduced with neutralized TCEP [5 mM, 30 min, room temperature(cat. no. 777205, Thermo Fisher Scientific)]; the native-free-thiol arm was held on ice without reduction. Both arms were then alkylated at free cysteines with the cysteine-reactive probe iodoacetamido-LC-phosphonic acid [CysPAT^66^, 10 mM, 1 h, room temperature, in the dark (cat. no. A52285, Thermo Fisher Scientific)], precipitated by chloroform–methanol, resuspended in 200 mM EPPS (pH 8.5), and digested with Trypsin/Lys-C (1:25, overnight, 37 °C).

Peptides (50 µg) were labeled with TMTpro 18-plex reagents (1:2.5 peptide:label, 1 h, room temperature). After ratio check, all channels were combined, desalted, enriched as described above for phosphopeptides, and analyzed by MS as described below. The flow-through was fractionated and analyzed as described below.

### Off-line basic pH reversed-phase (BPRP) fractionation (whole proteome)

We fractionated the pooled TMT-labeled peptides (specifically, the unbound and first wash from the phosphopeptides enrichment) *via* BPRP HPLC with an Agilent 1260 pump. Peptides were subjected to a 560 min linear gradient from 5% to 35% acetonitrile in 10 mM ammonium bicarbonate pH 8 at a flow rate of 0.25 mL/min over an Agilent 300Extend C18 column (3.5 μm particles, 2.1 mm ID, and 25 cm long). The peptide mixture was fractionated into a total of 96 fractions, which were consolidated into 24 super-fractions. Only a single set of 12 non-adjacent super-fractions were analyzed further. These fractions were subsequently acidified with 1% formic acid and vacuum centrifuged to near dryness. Each fraction was also desalted, dried by vacuum centrifugation, and reconstituted in 5% acetonitrile, 5% formic acid for LC–MS/MS.

### Liquid chromatography and mass spectrometry data acquisition for whole proteome analysis

Mass spectrometry data for 12 super-fractions were collected using a Orbitrap Exploris480 mass spectrometer (Thermo Fisher Scientific, San Jose, CA) coupled nLC-1200 liquid chromatograph. Peptides were separated on a 100 μm inner diameter microcapillary column packed with ∼35cm of Accucore C18 resin (2.6 μm, 150 Å, Thermo Fisher Scientific). For each analysis, we loaded ∼2 μg onto the column. Peptides were separated using a 120 min gradient of 2 to 40% acetonitrile in 0.125% formic acid with a flow rate of 300 nL/min. Spray voltage was set at 2700 V. The scan sequence began with an Orbitrap MS1 spectrum with the following parameters: resolution 60K, scan range 350-1350, automatic gain control (AGC) target 100%, maximum injection time 50ms, and centroid spectrum data type. We use a cycle time of 1s for MS2 analysis which consisted of HCD high-energy collision dissociation with the following parameters: resolution 50K, AGC 150%, maximum injection time 120ms, isolation window 0.7 Th, normalized collision energy (NCE) 35%, and centroid spectrum data type. Dynamic exclusion was set to 90sec. The FAIMS compensation voltages (CV) used were -35V/-45V/-60V for each set for 12 non-adjacent super-fractions.

### Liquid chromatography and tandem mass spectrometry for phosphoproteomics samples

Mass spectrometric data were collected on an Orbitrap Ascend MultiOmics mass spectrometer coupled to a Vanquish Neo UHPLC. Approximately 1µg of peptide was separated at a flow rate of 350 nL/min on a 100 µm capillary column that was packed with 35 cm of Accucore 150 resin (2.6 μm, 150Å; ThermoFisher Scientific). The gradient was 7-27% Buffer B (95% acetonitrile, 0.125% formic acid) which was mixed into buffer A (5% acetonitrile, 0.125% formic acid) over a 135 min gradient. Spray voltage was set at 2700 V. The scan sequence began with an MS1 spectrum (Orbitrap analysis, resolution 60,000, 350-1350 Th, automatic gain control (AGC) target is set to 125%, maximum injection time set to 25ms). The hrMS2 stage consisted of fragmentation by higher energy collisional dissociation (HCD, normalized collision energy 36%) and analysis using the Orbitrap (AGC 150%, maximum injection time 150 ms, isolation window 0.6 Th, resolution 45,000). The TopSpeed parameter was set at 1 second per CV. Data were acquired using three injections with three sets of compensation voltages (CVs): 40V/-60V/- 80V, -30V/-50V/-70V and -35/-45/-55V

#### Liquid chromatography and mass spectrometry data acquisition for free thiol fractional occupancy analysis

Mass spectrometry data were collected using a Orbitrap Astral mass spectrometer (Thermo Fisher Scientific, San Jose, CA) coupled with Neo Vanquish liquid chromatograph. We used the Astral detector of MS2 analysis. Peptides were separated on an Aurora gen 3 column (75 μm ID, 2 μm beads, 25 cm length (IonOpticks). For each analysis, we loaded ∼0.5 μg onto the column. Peptides were separated using a 135 min gradient of 6 to 27% acetonitrile in 0.125% formic acid with a flow rate of 300 nL/min. Spray voltage was set at 2700 V. The scan sequence began with an MS1 spectrum (Orbitrap analysis, resolution 60,000, 350-1350 Th, automatic gain control (AGC) target is set to 200%, maximum injection time set to 50ms). The hrMS2 stage consisted of fragmentation by higher energy collisional dissociation (HCD, normalized collision energy 35%) and analysis using the Astral mass analyzer (AGC 200%, maximum injection time 35ms, isolation window 0.7 Th). Dynamic exclusion was set to 20 s. Data were acquired using the FAIMSpro interface the dispersion voltage (DV) set to 5,000V. We used two injections with two different sets of compensation voltages, specifically -35V/-45V/-55V/-60V/-70V and -30/-40/-50V/- 65V.

#### Data analysis

Mass spectra were processed using a Comet-based in-house software pipeline. Spectra were converted to mzXML *via* MSconvert^67^. Database searching included all entries from the human UniProt reference database (downloaded: August 2021). The database was concatenated with one composed of all protein sequences for that database in the reversed order. Searches were performed using a 50-ppm precursor ion tolerance for total protein level profiling. The product ion tolerance was set to 0.03 Da. These wide mass tolerance windows were chosen to maximize sensitivity in conjunction with Comet searches and linear discriminant analysis^68,69^. TMTpro labels on lysine residues and peptide *N*-termini +304.207 Da), as well as carbamidomethylation of cysteine residues (+57.021 Da) were set as static modifications, while oxidation of methionine residues (+15.995 Da) was set as a variable modification. In addition, phosphorylation (+79.966 Da) at serine, threonine, and tyrosine residues were also set as variable modifications for phosphopeptide enrichment, while we used a variable modification of (+221.082 Da) on cysteine residues for CysPAT samples. Peptide-spectrum matches (PSMs) were adjusted to a 1% false discovery rate (FDR)^70,71^. PSM filtering was performed using a linear discriminant analysis, as described previously^69,71^, and then assembled further to a final protein-level FDR of 1%^71^. Proteins were quantified by summing reporter ion counts across all matching PSMs, as described previously^72^. For PTM site identification, the AScore10 false-discovery metric was used and only phosphosites that were “high-confidence”, with *P* ≤ 0.05, were retained. Reporter ion intensities were adjusted to correct for the isotopic impurities of the different TMTpro reagents according to manufacturer specifications. The signal-to-noise (S/N) measurements of peptides assigned to each protein were summed and these values were normalized so that the sum of the signal for all proteins in each channel was equal to account for differences in protein loading. Finally, each protein abundance measurement was scaled, such that the summed S/N for that protein across all channels equals 100, thereby generating a relative abundance measurement.

For global proteomics, summed, normalized reporter-ion abundances (8,066 quantified proteins) were log2-transformed, and differential protein abundance was tested with limma (v3.64.1; empirical-Bayes moderation with the trend and robust options) followed by DEqMS (v1.26.0), which replaces the prior variance with a peptide/PSM-count–dependent model, using pairwise contrasts against DMSO and BH adjustment; proteins were called significant at DEqMS-adjusted *P* < 0.05 and |log2 fold-change| ≥ log2(1.5). Pathway responses were assessed by pre-ranked GSEA (fgsea v1.34.2) over MSigDB collections on the DEqMS-ranked proteome. Analyses were performed in R v4.5.0.

For phosphoproteomics, normalized reporter-ion intensities were log2-transformed (4,229 quantified phosphosites). Differential phosphosite abundance was analyzed with limma-trend moderated t-statistics (eBayes, trend); sites were called significant at BH *q* < 0.05 and |log2FC| ≥ log2(1.5) (109 sites; 78 up, 31 down for OL2HG versus DMSO). Kinase activity was inferred from the phosphosite fold-changes using the Cantley Kinase Library motif atlas^73,74^ (kinase-library Python package v1.7.1): each 13-mer was scored against position-specific scoring matrices, the top 15 favored kinases were retained, and enrichment among regulated versus background sites was tested by one-sided Fisher’s exact test with BH adjustment; results were cross-checked with decoupleR^63^ (v2.14.0; ULM/KSEA over OmniPath). Phosphosite differential expression and the kinase activity inference used singly-phosphorylated sites only; the phosphoproteomics Supplementary Table 3 additionally reports multiply-phosphorylated (composite) sites, which are not included in Extended Data Fig. 5a–c or the reported site counts.

To resolve whether each phosphosite response required DGAT activity (Extended Data Fig. 5c), the four conditions were compared by DMSO-referenced and pairwise limma-trend contrasts [OL2HG *vs.* DMSO, OL2HG + DGATi *vs.* DMSO, DGATi *vs.* DMSO, OL2HG + DGATi *vs.* OL2HG, and OL2HG + DGATi *vs.* DGATi (each thresholded at BH *q* < 0.05 and |log₂ fold-change| ≥ log₂(1.5))]. Every quantified site was placed on a two-dimensional endpoint map, with x = log₂(OL2HG/DMSO) (the OL2HG effect) and y = log₂[(OL2HG + DGAT inhibitor)/DMSO] (the OL2HG effect on a DGAT-inhibited background), and assigned to exactly one category. Sites with a significant OL2HG-versus-DMSO effect were defined as OL2HG responders and sub-classified by the combination-versus-OL2HG contrast: DGAT-dependent when that contrast was significant and shifted the endpoint toward baseline (|y| < |x|, *i.e.,* DGAT inhibition reverses the response), DGAT-independent when it was non-significant (the response persists unchanged, lying on the y = x diagonal), or DGAT-amplified when it was significant and shifted the endpoint further from baseline (|y| > |x|). Sites lacking an OL2HG effect (|x| below the fold-change gate) but with a significant DGAT-inhibitor-versus-DMSO effect were defined as DGAT-inhibitor responders and sub-classified by the combination-versus-DGAT-inhibitor contrast into OL2HG-potentiated (significant and in the same direction as the DGAT- inhibitor effect) or OL2HG-independent (non-significant). Sites with neither a significant OL2HG nor DGAT-inhibitor effect but a significant combination-versus-DMSO effect were classified as combination-emergent (synergistic); all remaining sites were designated unchanged. Of the 4,229 quantified phosphosites, this partitioned 109 OL2HG responders (99 DGAT-independent, 7 DGAT-dependent, 3 DGAT-amplified), 10 DGAT-inhibitor- specific responders (9 OL2HG-independent, 1 OL2HG-potentiated) and 28 combination- emergent sites. Analyses were performed in R v4.5.0 (limma v3.64.1) and Python 3.

For redox proteomics, analysis was restricted to cysteines quantified in ≥2 PSMs with finite, positive reduced-thiol occupancy across all nine condition–replicate channel pairs (5,881 high-confidence sites). Because reduced-thiol occupancy is a bounded proportion with variance peaking near 50%, significance was assessed by limma-trend moderated t-statistics (eBayes, trend and robust; *n* = 3 per condition) computed on the variance-stabilized log₂(free/total) ratio, whereas the reported effect size was kept as the change in occupancy in percentage points (Δpp; positive = more reduced). A cysteine was defined as an OL2HG responder when it satisfied all three gates: *P* < 0.01, |Δpp| ≥ 10, excluding cysteines with non-physiological occupancy (*i.e.*, % reduced > 110% in either condition). This yielded 108 OL2HG responders (87 more oxidized, 21 more reduced). For each of these, DGAT-dependence was quantified by the reversal fraction, (mean_OL2HG − mean_OL2HG+DGATi)/(mean_OL2HG − mean_DMSO), *i.e.*, the share of the OL2HG redox shift that DGAT inhibition undoes (1 = the combination returns fully to the DMSO baseline; 0 = DGAT inhibition does nothing; < 0 = the shift overshoots further from baseline). Sites were classified as DGAT-dependent (reversal ≥ 0.5; the redox change is coupled to lipid storage; *n* = 56), DGAT-independent (−0.25 ≤ reversal < 0.5, or undefined; a direct, storage-uncoupled OL2HG effect; *n* = 50), or DGATi-amplified (reversal < −0.25; the shift is worsened by blocking storage; *n* = 2). Analyses were performed in R v4.5.0 (limma v3.64.1).

### Gene silencing by small interfering RNA (siRNA)

ON-TARGETplus siRNAs targeting human *HIF1A* (cat. no. L-004018-00-0005), *PLIN2* (cat. no. L-019204-01-0005), *SREBF1* (cat. no. L-006891-00-0005), and non-targeting control (cat. no. D-001810-10-05) were purchased from Dharmacon. HCMs were transfected with 5 pmol of each siRNA using Lipofectamine RNAiMAX transfection reagent (cat. no. 13778-150, Life Technologies). After transfection for 24 h, the cell medium was refreshed, and cells were then treated as reported in the results or Figure legends.

### Immunoblotting

Cells were lysed by sonication in RIPA lysis and extraction buffer (cat. no. 89900, Thermo Fisher Scientific) supplemented with Halt Protease Inhibitor Cocktail (cat. no. 78430, Thermo Fisher Scientific). Ten micrograms of total protein was loaded and separated on 4–15% pre-cast SDS–PAGE gels (cat. no. 4561086, Bio-Rad) using running buffer (cat. no. 161-0772, Bio-Rad) and then transferred to PVDF membranes using a semi-dry Turbo transfer system (cat. no. 170-4272, Bio-Rad). Protein blots were blocked with 5% (w/v) blotting grade milk (cat. no. 1706404XTU) prepared in Tris Buffered Saline with Tween (TBST, cat. no. 1706435 and 161-0781, Bio-Rad) and then incubated with specific primary antibodies overnight at 4 °C. HRP-linked secondary mouse or rabbit antibody (cat. no. 7076S and 7074S, Cell Signaling Technology; 1:2,000 dilution) and Immobilon detection reagents (cat. no. WBLUF0100, Sigma) were used to visualize protein blots. Images were captured using a ChemiDoc Touch Imaging system and quantitated by Image Lab software v.5.2 (Bio-Rad).

Antibodies used for immunoblotting were as follows: PLIN2 (15294-1-AP, Proteintech; 1:2,000), PLIN3 (10694-1-AP, Proteintech; 1:2,000), PLIN5 (26951-1-AP, Proteintech; 1:2,000), HIF-1α (610958, BD Biosciences; 1:500), β-actin–HRP (sc-47778 HRP, Santa Cruz Biotechnology; 1:1,000), SREBP1 (95879S, Cell Signaling Technology; 1:1,000), and SREBP2 (25940T, Cell Signaling Technology; 1:1,000).

### Mouse models

#### Animal housing ethics compliance

*L2hgdh^+/+^* and *L2hgdh*^−/−^ mice (aged-and sex-matched, 6–8-month-old) were maintained in the Brigham and Women’s Hospital animal facility under a 12 h light–dark cycle and controlled humidity (50–60%), with unrestricted access to water and standard chow (PicoDiet 5053, LabDiet; cat. no. 3005740-220). All experimental procedures were conducted in accordance with protocols approved by the Institutional Animal Care and Use Committee of Brigham and Women’s Hospital (2016N000573 and 2016N000241).

#### Baseline characterization

Male (*L2hgdh^+/+^*, *n* = 6; *L2hgdh*^−/−^, *n*= 5) and female (*L2hgdh^+/+^*, *n* = 6; *L2hgdh*^−/−^, *n*= 5) mice were euthanized by intraperitoneal overdose of ketamine (240–300 mg kg⁻¹) and xylazine (15– 30 mg kg⁻¹). Whole blood was immediately collected from the left ventricle into 2 mL tubes preloaded with heparin (10 µL; cat. no. NDC 25021-400-10), centrifuged (1,500 × *g*, 10 min, 4 °C), and plasma was flash-frozen in liquid nitrogen and stored at −80 °C until metabolomic and lipidomic analyses as described above. Hearts and livers were subsequently harvested, clamp-frozen in liquid N2, and stored at −80 °C until metabolomic and lipidomic analyses, as described above.

Cryopreserved livers and hearts obtained from male (*n* = 5) and female (*n* = 5) P11-old mice expressing wild-type (*n* = 5) or homozygous, global *L2hgdh* over-expression (*n* = 5), as reported previously^19^, were kindly provided by Prof. Chandel.

#### Langendorff perfusion

Hearts were isolated and perfused in Langendorff mode as described previously^30^. Briefly, mice were anticoagulated by intraperitoneal injection of 100 units of heparin 15 min before experiments and then euthanized with over-dose isoflurane (5% inhalation in isoflurane chamber). The hearts were quickly excised and arrested in ice-cold regular Krebs-Henseleit (KH) buffer containing (in mM) NaCl 118, KCl 5.3, CaCl2 2.5, MgSO4 1.2, EDTA 0.5, NaHCO3 25, glucose 10, prepared freshly and equilibrated with 95% O2 / 5% CO2, yielding a pH of 7.40, and connected *via* the aorta to the perfusion cannula. Right ventricular drainage was accomplished by incision of the pulmonary artery. The effluent from the thebesian veins was drained by a thin polyethylene tube (PE-10) pierced through the apex of the left ventricle. For baseline perfusion of male hearts (*L2hgdh^+/+^*, *n* = 5; *L2hgdh*^−/−^, *n* = 7), isolated hearts were perfused in the Langendorff mode at a constant coronary perfusion pressure of 80 mmHg at 37 °C with the modified KH buffer as mentioned above for 30 min to equilibrate followed by an additional 30 min of base-line perfusion. For low-flow ischemia of male hearts (*L2hgdh^+/+^*, *n* = 8; *L2hgdh*^−/−^, *n* = 5), hearts were perfused as per the baseline-perfusion protocol, with the addition of 90 min perfusion at 10 mmHg. These perfusion protocols are adapted directly from our previous publication^30^.

Acylcarnitine responses in Langendorff-perfused hearts were modeled by two-way fixed-effects ANOVA (genotype × perfusion condition) on log2 peak area, reporting the genotype, condition, and interaction terms. Analyses were performed in R v4.5.0.

### Bioinformatic re-analysis of published datasets

Associations between circulating 2HG and triglycerides in humans were examined in three previously published^52^ plasma metabolomic lifestyle/weight-loss cohorts from three weight-loss intervention cohorts (The Weight Loss Maintenance, WLM^53^, NCT00054925; The Studies of Targeted Risk Reduction Interventions through Defined Exercise in individuals with Pre-Diabetes, STRRIDE-PD^54,55^, NCT00962962; and The Columbia Bariatric and Diabetes, CBD^56–60^, NCT01516320, NCT02287285, NCT02929212, NCT00571220) and is openly available at Zenodo (https://doi.org/10.5281/zenodo.4767969). Within each cohort, 2HG was correlated with each triglyceride species (Spearman), and the dependence of the 2HG–triglyceride correlation on acyl-chain carbon number was summarized across species and compared between cohorts using the Fisher z-transformation.

The *L2hgdh*-overexpression RNA-seq, re-analyzed in Extended Data Fig. 9n from a dataset published in Ref.^19^, was reprocessed through the DESeq2/fgsea pipeline described above. Analyses used R v4.5.0 and Python 3 (SciPy v1.15.3, statsmodels v0.14.6).

### Statistics

The transcriptomic, proteomic, phosphoproteomic, cysteine-redox, lipidomic, metabolomic and stable-isotope-flux datasets were analyzed in R (v4.5.0; Bioconductor 3.21) and Python 3 as detailed in the corresponding sections, with multiple-hypothesis testing controlled by the Benjamini–Hochberg false-discovery rate. Principal software comprised limma (v3.64.1), DEqMS (v1.26.0), DESeq2 (v1.48.2), fgsea (v1.34.2), decoupleR (v2.14.0), emmeans (v2.0.2), apeglm (v1.30.0) and ggplot2 (v4.0.2) in R, and kinase-library (v1.7.1), SciPy (v1.15.3) and statsmodels (v0.14.6) in Python.

All other statistical analyses were performed using GraphPad Prism (version 9.0). Data normality was assessed using the Shapiro–Wilk test. For datasets that satisfied normality assumptions, statistical significance among three or more groups was evaluated using one-way ANOVA, followed by either Tukey’s or Dunnett’s post hoc multiple-comparison test, as appropriate. Comparisons between two independent groups were performed using an unpaired two-tailed Student’s t-test. For datasets that did not meet normality assumptions, statistical significance among three or more groups was assessed using the Kruskal–Wallis test, followed by Dunn’s post hoc test, while comparisons between two experimental conditions were conducted using the non-parametric Mann–Whitney U test. Data collection and analysis were performed blinded to experimental conditions whenever feasible. All data are presented as mean ± standard error of mean (SEM).

---

**Extended Data Fig. 1:**
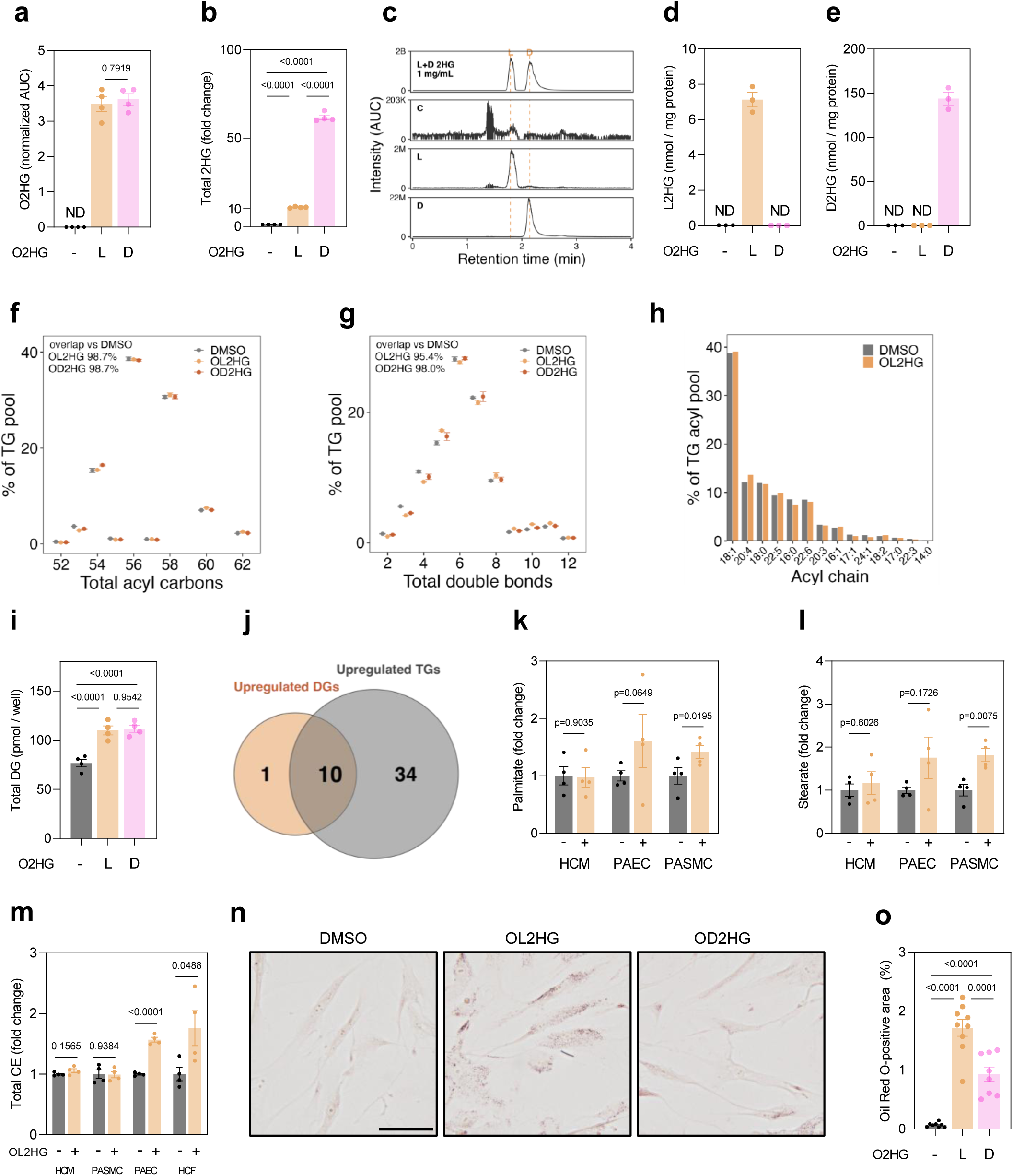
Stereospecific 2HG delivery and proportional TG expansion. **a**–**e,** Total intracellular octyl-2HG (O2HG) (**a**), free total 2HG (**b**), and stereoselective L2HG and D2HG (**c**–**e**) abundance in HCMs treated with DMSO, OL2HG (500 µM), or OD2HG (500 µM) for 24 h; *n* = 3–4 per group. **f,g,** Composition of the OL2HG-responsive TG pool, comprising 39 species, across total acyl carbon number (**f**) and total double bond number (**g**) in HCMs treated with DMSO, OL2HG (500 µM), or OD2HG (500 µM) for 24 h. Each bin is plotted as its percentage share of that group’s total TG signal. Overlap percentages indicate the shared area between distributions. *n* = 4 per group. **h,** TG acyl-chain composition in HCMs treated with DMSO or OL2HG (500 µM) for 24 h. **i,** Total DG abundance in HCMs treated with DMSO, OL2HG (500 µM), or OD2HG (500 µM) for 24 h; *n* = 4 per group. **j,** Overlap between acyl-chain compositions of OL2HG-increased DG and OL2HG-increased TG species. Significance was defined as BH-adjusted *P* < 0.05 and ≥1.5-fold change. **k,l,** Free palmitate (**k**) and stearate (**l**) abundance in HCMs, PAECs, and PASMCs treated with DMSO or OL2HG (500 µM) for 24 h; *n* = 4 per group per cell type. **m,** Total cholesteryl ester (CE) abundance in HCMs, PASMCs, PAECs and HCFs treated with DMSO or OL2HG (500 µM) for 24 h; *n* = 4 per group per cell type. **n,o,** Representative 40**×** images (**n**) and quantification (**o**) of Oil-Red-O staining of neutral lipids in PASMCs treated with DMSO, OL2HG (500 µM), or OD2HG (500 µM) for 24 h. Scale bar, 200 µm. Unless otherwise stated, data are mean ± s.e.m. For two-group comparisons, two-tailed unpaired t-tests or Mann–Whitney tests were used; for multiple-group comparisons, one-way ANOVA with Tukey’s test or Kruskal–Wallis tests with Dunn’s test were used, according to data distribution. Exact *P* values are shown.

**Extended Data Fig. 2:**
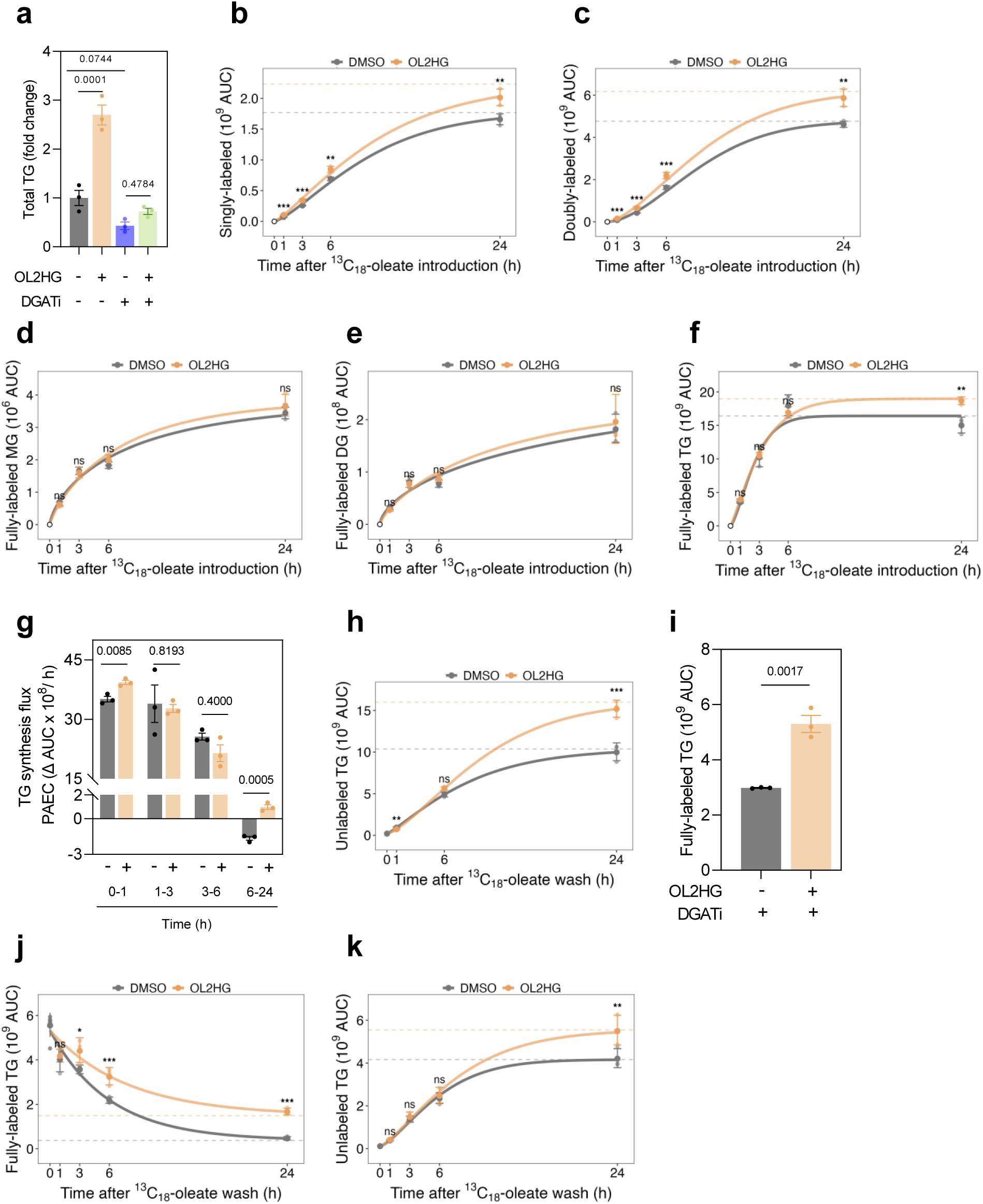
Stable-isotope tracing localizes L2HG-induced TG synthesis to the terminal DGAT step. **a,** Total TG abundance in HCMs treated with DMSO, OL2HG (500 µM), DGATi (A- 922500 and PF-06424439, 10 µM each), or OL2HG + DGATi for 24 h; *n* = 3 per group. **b–e,** Absolute abundance of singly labeled triolein [TG(1 × ^13^C18-18:1; 2 × ^12^C18-18:1)] (**b**), doubly labeled triolein [TG(2 × ^13^C18-18:1; 1 × ^12^C18-18:1)] (**c**), fully labeled monoolein [^13^C18-MG(18:1)] (**d**), and fully labeled diolein [DG(2 × ^13^C18-18:1)] (**e**) during the [U-¹³C₁₈]-oleate (50 µM) pulse in HCMs pre-treated with DMSO or OL2HG (500 µM) for 24 h. **f,g** Absolute abundance (**f**) and flux (**g**) of fully labeled triolein during the [U- ¹³C₁₈]-oleate (50 µM) pulse in PAECs pre-treated with DMSO or OL2HG (500 µM) for 24 h. **h,** Absolute abundance of unlabeled triolein in HCMs pulsed with [U-¹³C₁₈]-oleate (50 µM) for 24 h and chased with DMSO or OL2HG (500 µM). **i,** Fully labeled triolein remaining in HCMs pulsed with [U-¹³C₁₈]-oleate (50 µM) for 24 h and chased with DGATi (A-922500 and PF-06424439, 10 µM each) ± OL2HG (500 µM) for 24 h. **j,k,** Clearance of fully labeled triolein (**j**) and appearance of unlabeled triolein (**k**) in HCFs pulsed with [U-¹³C₁₈]-oleate (50 µM) for 24 h and chased with DMSO or OL2HG (500 µM). All flux curves show through-origin Weibull fits (pulse design) or exponential-to-plateau fits (pulse–chase design); dashed lines indicate fitted plateaus; points show geometric mean ± geometric s.d.; *n* = 3 per condition per time point, except chase t = 0 where *n* = 6. *P* values were calculated by two-way ANOVA with Holm-adjusted OL2HG–DMSO contrasts within each time point. For all other panels, data are mean ± s.e.m. For two-group comparisons, two-tailed unpaired t-tests or Mann–Whitney tests were used; for multiple-group comparisons, one-way ANOVA with Tukey’s test or Kruskal–Wallis tests with Dunn’s test were used, according to data distribution. Exact *P* values are shown.

**Extended Data Fig. 3:**
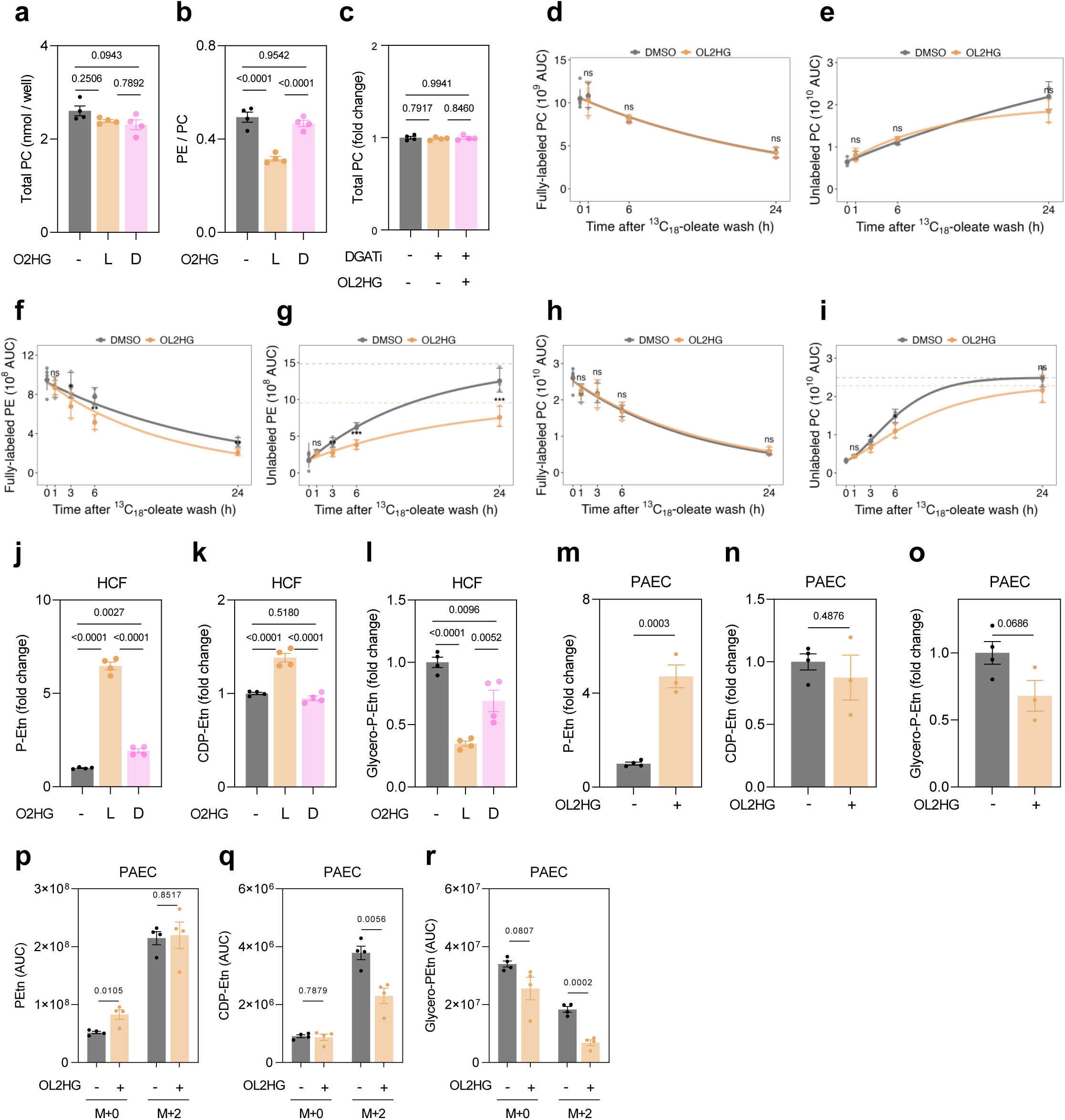
L2HG selectively constrains synthesis of PE, not PC, across cell types. **a,** Total PC abundance (**a**) and PE-to-PC ratio (**b**) in HCMs treated with DMSO, OL2HG (500 µM), or OD2HG (500 µM) for 24 h; *n* = 4 per group. **c,** Total PC abundance in PAECs treated with DGATi (A-922500 and PF-06424439, 10 µM each) ± OL2HG (500 µM) for 24 h; *n* = 4 per group. **d,e,** Clearance of fully labeled dioleoyl-PC [PC(2 × ^13^C18-18:1)] (**d**) and appearance of unlabeled dioleoyl-PC [PC(2 × ^12^C18-18:1)] (**e**) in HCMs pulsed with [U-¹³C₁₈]-oleate (50 µM) for 24 h and chased with DMSO or OL2HG (500 µM). **f–i,** Clearance of fully labeled dioleoyl-PE [PE(2 × ^13^C18-18:1)] (**f**) and appearance of unlabeled dioleoyl-PE [PE(2 × ^12^C18-18:1)] (**g**), as well as fully labeled (**h**) and unlabeled (**i**) dioleoyl-PC, in HCFs pulsed with [U-¹³C₁₈]-oleate (50 µM) for 24 h and chased with DMSO or OL2HG (500 µM). **j–o,** Ethanolamine-branch Kennedy-pathway intermediates in HCFs (**j**–**l**) and PAEC (**m**–**o**) following treatment with DMSO, OL2HG (500 µM), or OD2HG (500 µM) for 24 h. *n* = 3–4 per group. **p–r,** [^13^C2]-ethanolamine labeling of P-Etn (**p**), CDP-Etn (**q**), and glycero-P-Etn (**r**) in PAECs treated with DMSO or OL2HG (500 µM) for 24 h, showing the naturally abundant (M+0) and the fully labeled (M+2) isotopologues. *n* = 4 per group. All flux curves show through-origin Weibull fits (pulse design) or exponential-to-plateau fits (pulse–chase design); dashed lines indicate fitted plateaus; points show geometric mean ± geometric s.d.; *n* = 3 per condition per time point, except chase t = 0 where *n* = 6. *P* values were calculated by two-way ANOVA with Holm-adjusted OL2HG– DMSO contrasts within each time point. For all other panels, data are mean ± s.e.m. For two-group comparisons, two-tailed unpaired t-tests or Mann–Whitney tests were used; for multiple-group comparisons, one-way ANOVA with Tukey’s test or Kruskal–Wallis tests with Dunn’s test were used, according to data distribution. Exact *P* values are shown.

**Extended Data Fig. 4:**
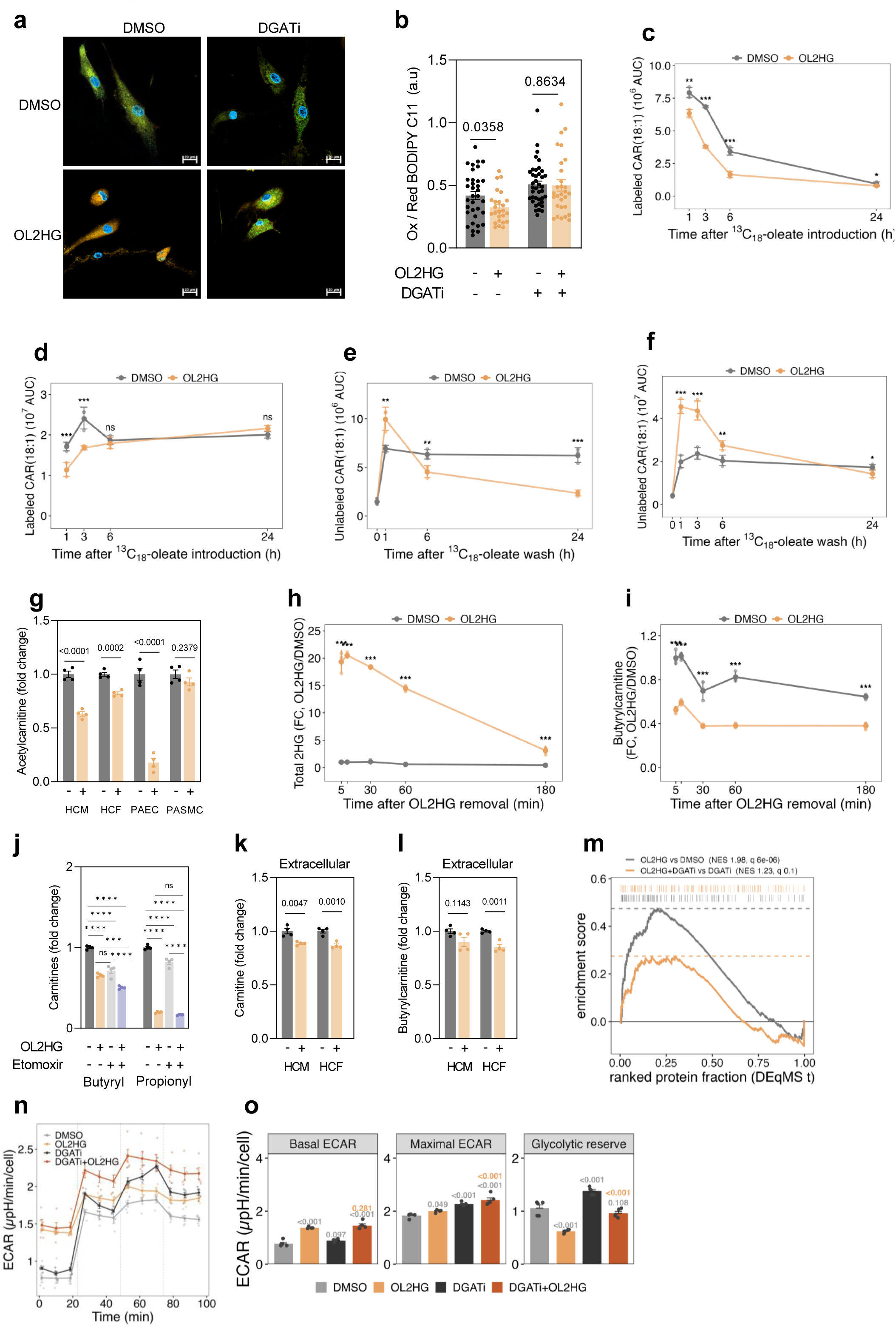
Lipid peroxidation, acylcarnitine dynamics, and glycolytic shift downstream of L2HG. **a,b,** Representative 63**×** BODIPY-C11 oxidized (green) / reduced (orange) images (**a**) and oxidized/reduced ratio quantification (**b**) in oleate (50 µM)-loaded HCMs treated with DMSO, OL2HG (500 µM), DGATi (A-922500 and PF-06424439, 10 µM each), or OL2HG + DGATi for 24 h. Scale bar, 20 µm. *n* = 25–39 cells from four independent samples per group. **c,d,** Labeled CAR(18:1) abundance in PAECs (**c**) and HCFs (**d**) pre-treated with DMSO or OL2HG (500 µM) for 24 h and then pulsed with [U-¹³C₁₈]-oleate (50 µM). **e,f,** Unlabeled ^13^C18-oleoylcarnitine [CAR(18:1)] abundance in HCMs (**e**) and HCFs (**f**) pulsed with [U-¹³C₁₈]-oleate (50 µM) for 24 h and chased with DMSO or OL2HG (500 µM). **g,** Acetylcarnitine abundance in HCMs, HCFs, PAECs, and PASMCs treated with DMSO or OL2HG (500 µM) for 24 h. **h,i,** Total intracellular 2HG (**h**) and butyrylcarnitine (**i**) in HCMs after OL2HG pre-treatment for 24 h and media wash. **j,** Butyrylcarnitine and propionylcarnitine abundance in HCMs treated with OL2HG (500 µM) ± etomoxir (10 µM) for 24 h. **k,l,** Extracellular free carnitine (**k**) and butyrylcarnitine (**l**) in HCM and HCF medium following treatment with DMSO or OL2HG (500 µM) for 24 h. **m,** Running enrichment plot for fatty acid metabolism hallmark gene set ranked by the proteome DEqMS t-statistic for OL2HG *vs* DMSO (grey) and OL2HG + DGATi *vs* DGATi (orange) using fgsea. Normalized enrichment score (NES) and *q* values are shown. **n,o,** Seahorse ECAR kinetics (**n**) and parameters (**o**) in HCMs treated with DMSO, OL2HG, DGATi or OL2HG + DGATi for 24 h; *n* = 4–5 per group. For flux curves (**c– f,h,i**), points show geometric mean ± geometric s.d.; *n* = 3 per condition per time point, except chase t = 0 where *n* = 6. *P* values were calculated by two-way ANOVA with Holm-adjusted OL2HG–DMSO contrasts within each time point. For all other panels, data are mean ± s.e.m. For two-group comparisons, two-tailed unpaired t-tests or Mann–Whitney tests were used; for multiple-group comparisons, one-way ANOVA with Tukey’s test or Kruskal–Wallis tests with Dunn’s test were used, according to data distribution. Exact *P* values are shown.

**Extended Data Fig. 5.**
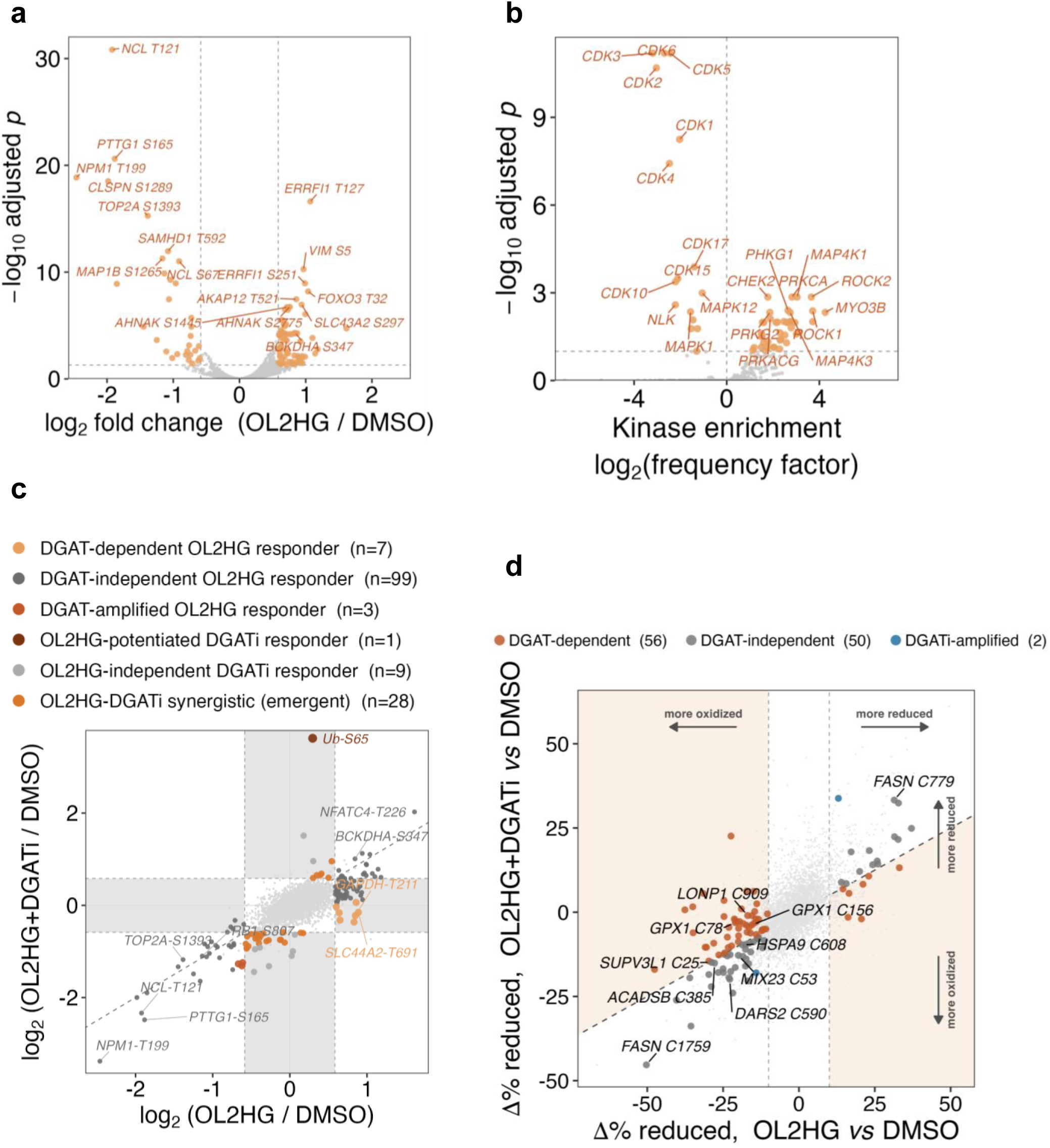
OL2HG-dependent changes to the phospho- and redox- proteomes. **a,** Volcano plot of TMT phosphoproteomics (OL2HG *vs* DMSO); 109 sites changed (78 up, 31 down; limma-trend moderated t, BH *P* < 0.05 and |log2FC| ≥ log21.5; *n* = 3 per group). **b,** Kinase-activity enrichment (Cantley Kinase Library motif atlas; one-sided Fisher exact, BH *P* < 0.10) for regulated phosphosites; x, signed log2 frequency factor (positive, activated); proline-directed CDK/ERK kinases suppressed, basophilic AGC kinases activated. **c,** Phosphosite DGAT-dependence: log2FC(OL2HG + DGATi / DMSO) *vs* log2FC(OL2HG / DMSO); 99 of 109 OL2HG-responsive sites were DGAT- independent. Dashed line, y = x. Representative sites are labeled. **d,** Cysteine-redox DGAT-dependence (free thiol stoichiometry, % reduced = −TCEP/+TCEP): Δ% reduced (OL2HG + DGATi / DMSO) *vs* Δ% reduced (OL2HG / DMSO) for 108 OL2HG-responsive cysteines of 5,881 quantified (|Δ| ≥ 10 percentage points and *P* < 0.01; *n* = 3 per group); 56 DGAT-dependent, 50 DGAT-independent, 2 DGAT-amplified. Dashed line, reversal = 0.5 boundary. Representative sites are labeled.

**Extended Data Fig. 6:**
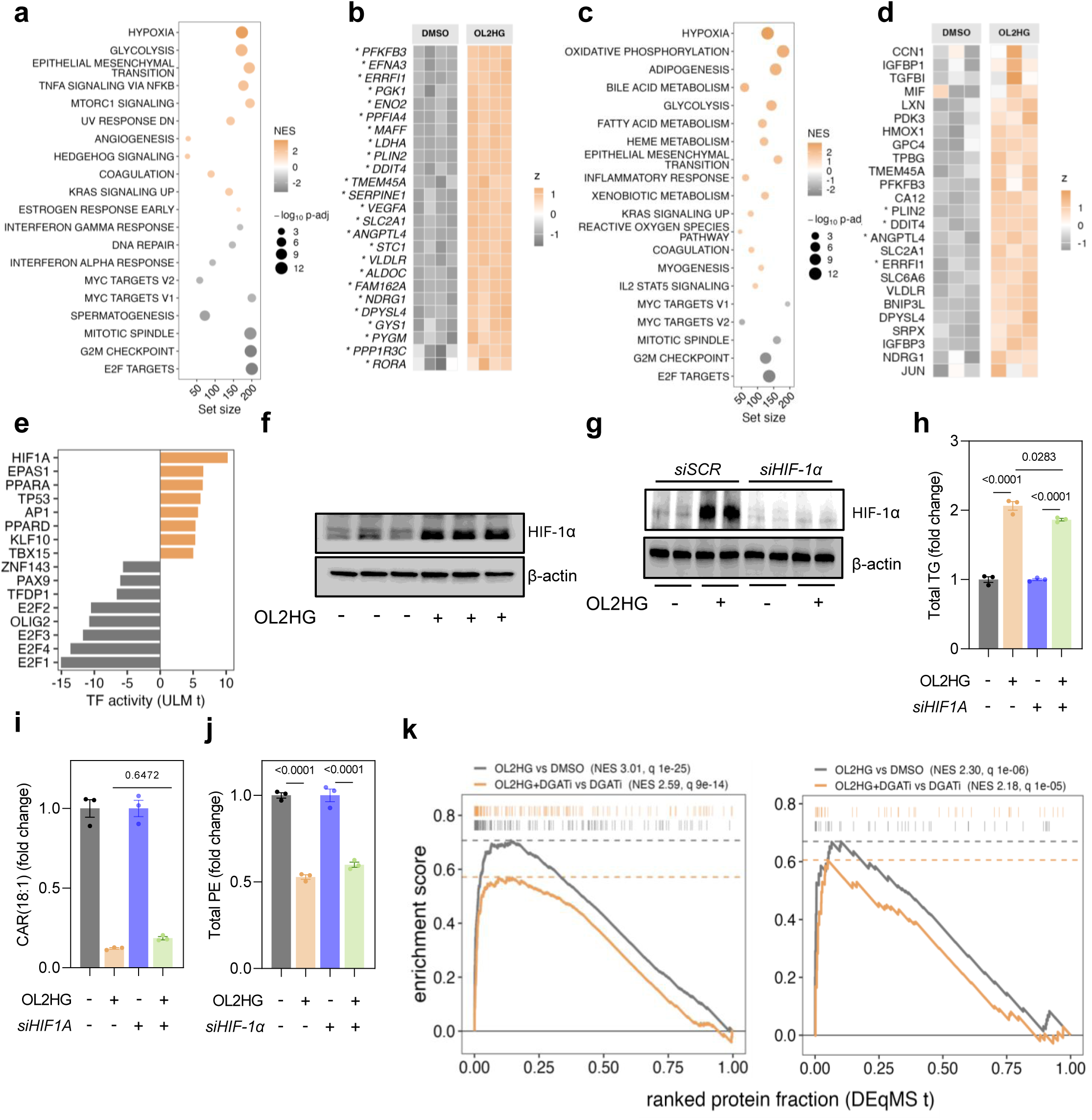
L2HG-induced pseudo-hypoxic transcription is dispensable for lipid rewiring. **a,** Hallmark gene-set enrichment of the HCM transcriptome after OL2HG treatment (500 µM) *vs* DMSO for 24 h. Genes were ranked by DESeq2 Wald statistic; *n* = 4 per group. **b,** Leading-edge heatmap of hypoxia genes from **a**, shown as per-gene *z*-scores of variance-stabilized counts. **c,d,** Hallmark gene-set enrichment (**c**) and hypoxia leading-edge heatmap (**d**) of the HCM proteome after OL2HG treatment (500 µM) *vs* DMSO for 24 h. Proteins were ranked by DEqMS t-statistic; *n* = 3 per group. **e,** Transcription factor activity inferred by decoupleR univariate linear model using CollecTRI regulons and DESeq2 Wald statistics as input. **f,** HIF-1α immunoblot in HCMs ± OL2HG (500 µM, 24 h); β-actin, loading control. **g,** *HIF1A* knockdown validation immunoblot in HCMs treated with *siSCR* or *siHIF1A* ± OL2HG (500 µM, 24 h). **h–j,** Total TG (**h**), CAR(18:1) (**i**), and total PE (**j**) in HCMs treated with *siSCR* or *siHIF1A* ± OL2HG (500 µM, 24 h). Panel **h** is normalized within each siRNA condition: OL2HG-treated *siSCR* cells are plotted relative to DMSO-treated *siSCR* cells and OL2HG-treated *siHIF1A* cells relative to DMSO-treated *siHIF1A* cells. **k,** Proteomic running enrichment of hypoxia hallmark (left) and HIF1 pathway (right) signatures in HCMs treated with OL2HG (500 µM) ± DGATi (A-922500 and PF-06424439, 10 µM each) for 24 h. Unless otherwise stated, data are mean ± s.e.m. For two-group comparisons, two-tailed unpaired t-tests or Mann–Whitney tests were used; for multiple-group comparisons, one-way ANOVA with Tukey’s test or Kruskal–Wallis tests with Dunn’s test were used, according to data distribution. Exact *P* values are shown. NES, normalized enrichment score.

**Extended Data Fig. 7:**
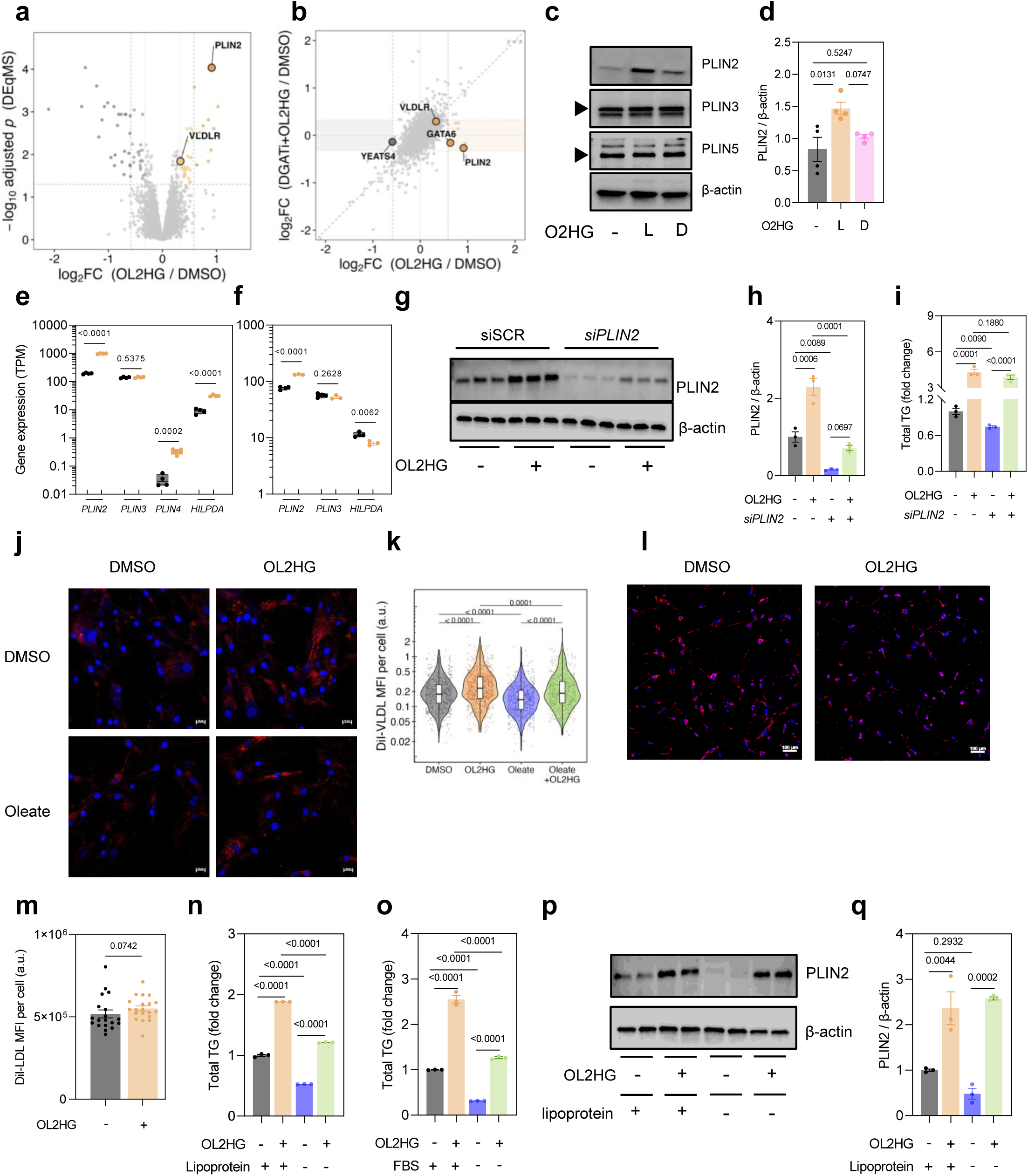
L2HG-induced PLIN2 and VLDLR are dispensable for TG accumulation. **a,** Volcano plot of global proteomics in HCMs treated with OL2HG (500 µM, 24 h) *vs* DMSO. Proteins were analysed by limma and DEqMS with BH adjustment; *n* = 3 per group. **b,** DGAT-dependence of the OL2HG-regulated proteome, shown as log₂FC(OL2HG + DGATi / DMSO) *vs* log₂FC(OL2HG / DMSO). Dashed line, y = x; shaded bands indicate reversal toward baseline. **c,d,** PLIN2, PLIN3, and PLIN5 immunoblots (**c**) and PLIN2 quantification (**d**) in HCMs treated with DMSO, OL2HG (500 µM), or OD2HG (500 µM) for 24 h. **e,f,** Lipid droplet-associated transcript abundance in HCMs (**e**) and PAECs (**f**) treated with DMSO or OL2HG (500 µM) for 24 h, shown as transcripts per million (TPM). **g–i,** *siPLIN2* validation immunoblot (**g**), PLIN2 quantification (**h**), and total intracellular TG (**i**) in HCMs ± OL2HG (500 µM, 24 h). **j,k,** Confocal microscopy representative 40**×** images (**j**) and quantification (**k**) of Dil-VLDL in HCMs treated with OL2HG (500 µM) ± oleate (50 µM) for 24 h. Nuclei are stained with Hoechst. Scale bar, 20 µm. *n* = 506–807 cells from four independent experiments. **l,m,** Confocal microscopy representative 10**×** images (**l**) and quantification (**m**) of DiI-LDL in HCMs treated with DMSO or OL2HG (500 µM) for 24 h. Nuclei are stained with Hoechst. Scale bar, 100 µm; *n* = 19–20 fields each containing 72–210 cells from four independent experiments. **n,o,** Total TG abundance in HCMs ± OL2HG (500 µM, 24 h) in lipoprotein- free medium (**n**) or serum-free medium (**o**). *n* = 3 per group. **p,q,**. PLIN2 immunoblot (**p**) and quantification (**q**) in HCMs ± OL2HG (500 µM, 24 h) in the presence or absence of lipoprotein-free medium. Unless stated otherwise, data are mean ± s.e.m. For two-group comparisons, two-tailed unpaired t-tests or Mann–Whitney tests were used; for multiple-group comparisons, one-way ANOVA with Tukey’s test or Kruskal–Wallis tests with Dunn’s test were used, according to data distribution. Exact *P* values are shown.

**Extended Data Fig. 8:**
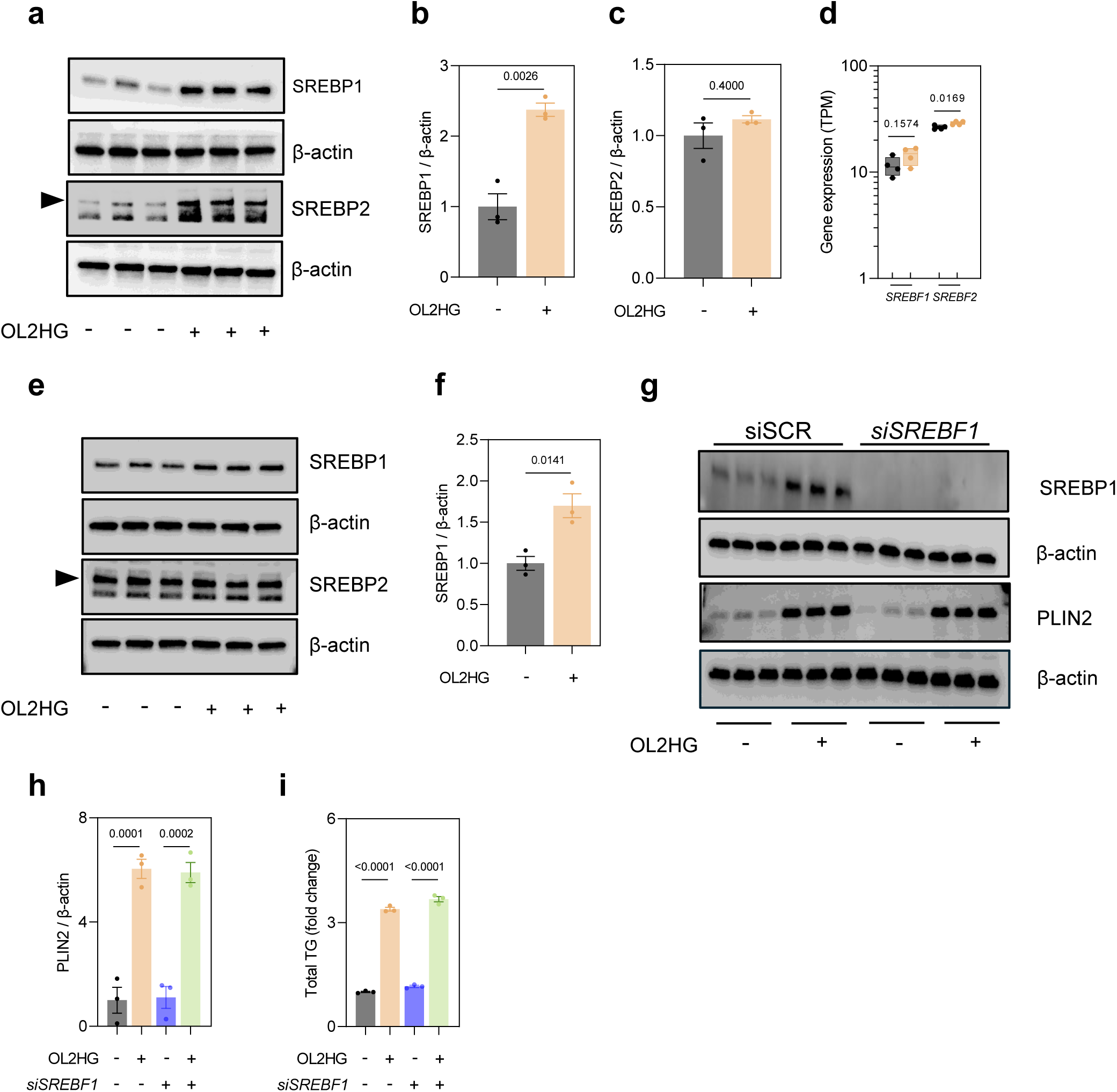
L2HG-induced SREBP1 stabilization is dispensable for TG accumulation. **a–c,** SREBP1 and SREBP2 immunoblot (**a**) and quantification (**b**–**c**) in HCMs treated with DMSO or OL2HG (500 µM) for 24 h. **d,** *SREBF1* and *SREBF2* transcript abundance in HCMs treated with DMSO or OL2HG (500 µM) for 24 h, shown as transcripts per million (TPM). **e,f,** SREBP1 and SREBP2 immunoblot (**e**) and quantification (**f**) in PAECs treated with DMSO or OL2HG (500 µM) for 24 h. **g,** Immunoblot of SREBP1 and PLIN2 in HCMs treated with *siSREBF1* and OL2HG (500 µM) for 24 h. **h,i,** PLIN2 immunoblot quantification (**h**) and total TG abundance (**i**) in HCMs treated with *siSCR* or *siSREBF1* ± OL2HG (500 µM) for 24 h. For all panels, data are mean ± s.e.m. For two-group comparisons, two-tailed unpaired t-tests or Mann–Whitney tests were used; for multiple-group comparisons, one-way ANOVA with Tukey’s test or Kruskal–Wallis tests with Dunn’s test were used, according to data distribution. Exact *P* values are shown.

**Extended Data Fig. 9:**
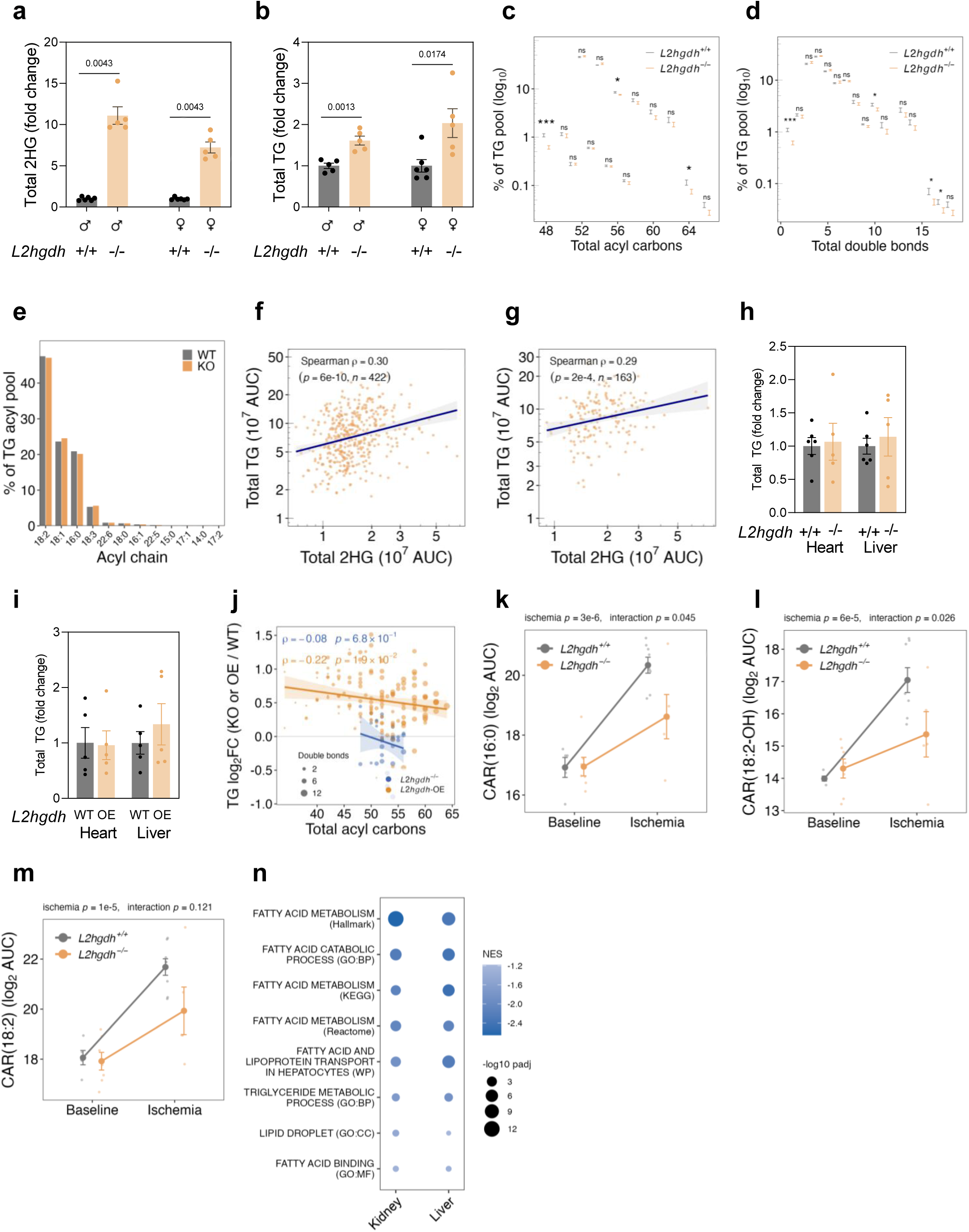
*In vivo* 2HG accumulation is associated with circulating TGs and decreased ischemic acylcarnitines. **a,b,** Plasma total 2HG (**a**) and total TG (**b**) in *L2hgdh^−/−^* (*n* = 5 per sex) *vs. L2hgdh^+/+^* (*n* = 5–6 per sex) mice by sex. **c,d,** Distribution of the plasma TG pool across total acyl carbon number (**c**) and total double bond number (**d**) in *L2hgdh^−/−^* (*n* = 5 per sex) *vs. L2hgdh^+/+^* (*n* = 6 per sex) mice. All 60 detected TG species are shown, with sexes pooled and each bin expressed as the percentage share of that genotype’s total TG signal. **e,** TG acyl chain composition in plasma from *L2hgdh^−/−^*(*n* = 5 per sex) *vs. L2hgdh^+/+^* (*n* = 6 per sex) mice. **f,g** Circulating 2HG *vs.* the shorter-chain (≤ 50 acyl-carbon) TG pool in two human cohorts: WLM (**f**; *n* = 422) and STRRIDE-PD (**g**; *n* = 163). **h,i,**. Heart and liver total TG abundance in male *L2hgdh^−/−^* (*n* = 5) *vs. L2hgdh^+/+^* (*n* = 6) mice (**h**) and in male and female *L2hgdh*-overexpressing mice (*n* =5 per genotype) (**i**). **j,** Per-species TG fold change plotted against total acyl chain carbon number in *L2hgdh^−/−^*(blue, *n* = 5–6 per genotype) and *L2hgdh*- overexpressing (orange, *n* = 5 per genotype) livers *vs.* their respective wild-type control animals. Solid lines show linear regression with 95% confidence intervals. **k–m,** Individual long-chain acylcarnitines in Langendorff-perfused hearts at baseline *vs.* ischemia in *L2hgdh^−/−^ vs L2hgdh^+/+^* mice: CAR(16:0) (**k**), CAR(18:2-OH) (**l**), and CAR(18:2) (**m)**. *n* = 5–8 hearts per group analyzed by a two-way ANOVA. **n,** Gene-set enrichment of fatty acid and TG metabolism programs of RNA-seq data from kidney and liver in *L2hgdh*- overexpressig mice. Genes were ranked by DESeq2 Wald statistic; *n* = 5 per group per tissue. Unless otherwise stated, data are mean ± s.e.m.; for two-group comparisons, two-tailed unpaired t-tests or Mann–Whitney tests were used, according to data distribution. Exact *P* values shown. Panels **f**,**g**, and **n** comprise data from publicly available datasets (see Data availability).

